# mRNA-LNP therapy restores systemic nucleoside imbalance in a mitochondrial DNA depletion syndrome

**DOI:** 10.64898/2026.09.08.750057

**Authors:** Nissa L Carrodus, Jenny J Yang, Javier Ramón, Keira Turner, Yamini M Kanse, Stavroula Petridi, Zitian Xiao, Abhilesh Dhawanjewar, Theodore E Leonard, George Nolan, Annelies Quaegebeur, Ramon Martí, Seth W Cheetham, Jelle van den Ameele

## Abstract

Mitochondrial DNA depletion syndromes (MDS) are inherited conditions caused by pathogenic variants in mitochondrial DNA maintenance genes. Most MDS are severe, fatal and incurable conditions. Mitochondrial neurogastrointestinal encephalomyopathy (MNGIE) is an MDS resulting from loss-of-function mutations in the *TYMP* gene, encoding Thymidine Phosphorylase (TP). Systemic nucleoside accumulation caused by TP deficiency disrupts mitochondrial nucleotide homeostasis, which underlies disease progression and mortality. Current treatments, including liver and hematopoietic stem cell transplantation, partially restore TP activity but are invasive, carry substantial risk and are limited by donor availability. Here, we establish *TYMP*-mRNA in lipid nanoparticles (*hTYMP*-mRNA-LNPs) as a safe non-viral protein-replacement therapy for MNGIE. Intravenous administration of *hTYMP*-mRNA-LNPs induced robust hepatic TP expression in a mouse model of MNGIE, was well tolerated and restored circulating nucleosides to wild-type levels within hours, lasting up to three weeks, at a preclinical minimally effective dose of 0.25mg/kg. To facilitate repeat administration and patient access, we demonstrate enhanced efficiency of subcutaneous mRNA-LNP delivery by co-administration of recombinant or mRNA-encoded (*SPAM1*-mRNA-LNPs) hyaluronidase, achieving effective hepatic TP expression and systemic nucleoside clearance. These findings establish mRNA-LNP-mediated protein replacement as a therapeutic strategy for a primary mitochondrial disease where transient liver-targeted expression is sufficient to correct a systemic metabolic defect. More broadly, our results support the development of mRNA-LNP therapeutics and their subcutaneous delivery as a generalizable platform for treating monogenic diseases, through repeatable, non-viral protein replacement.

## INTRODUCTION

Mitochondrial DNA depletion syndromes (MDS) are a group of rare diseases caused by inherited pathogenic variants in nuclear genes that control mitochondrial DNA (mtDNA) replication, maintenance or nucleotide metabolism (*1, 2*). Clinical presentation is highly diverse, and can range from infantile-onset fatal multi-system disease to late-onset isolated eye muscle weakness. Recent progress in therapeutic development (*3*), including the approval of deoxynucleotide supplementation as a disease-modifying treatment for thymidine kinase 2 (TK2) deficiency (*4*), opens new prospects for these often-fatal conditions. However, most remain without cure. Development of safe and effective gene therapies for mitochondrial disease was recently identified as one of the top ten research priorities by a Structured Priority-Setting-Partnership between mitochondrial disease patients, families and health care providers (*5*), underscoring the importance of addressing this high unmet clinical need, but with important barriers to clinical translation (*6*).

Mitochondrial neurogastrointestinal encephalomyopathy (MNGIE), is an autosomal recessive MDS caused by loss-of-function variants in *TYMP*, the gene encoding the enzyme thymidine phosphorylase (TP). Loss of TP function leads to systemic elevation of thymidine and deoxyuridine nucleosides (dThd/dUrd) and imbalance of the mitochondrial deoxynucleoside triphosphate (dNTP) pool, causing progressive mtDNA maintenance defects (*7–10*). The clinical phenotype of MNGIE results from the disruption of mitochondrial oxidative phosphorylation due to accumulated mtDNA mutations and mtDNA depletion. MNGIE symptom onset is typically in the second decade of life, and is rapidly progressive after diagnosis, with the majority of patients not surviving beyond 40 years of age (*11, 12*). The most debilitating symptom of MNGIE is gastrointestinal (GI) dysmotility, which occurs in almost all reported patients. Other typical symptoms include cachexia, chronic progressive eye muscle weakness and peripheral neuropathy, and all patients have asymptomatic leukoencephalopathy detectable on brain MRI (*13*).

MNGIE disease severity correlates with residual TP activity. Heterozygous mutation-carriers remain completely asymptomatic despite relatively low TP activity (∼25-35% of the normal value) (*14*), and late-onset MNGIE patients with symptom-onset in 40s-50s have some residual TP activity (∼10-15% of healthy controls), causing a more modest accumulation of dThd/dUrd (*15*). Thus, plasma dThd/dUrd levels constitute effective biomarkers for diagnosis and monitoring of treatment efficacy (*16*).

Several treatment strategies have been proposed for MNGIE that are aimed at restoring systemic nucleoside balance, either by directly clearing dThd/dUrd from the circulation, or by reconstituting TP activity in part of the body (*13, 14, 17, 18*). Because unphosphorylated nucleosides are readily transported across all plasma membranes through dedicated transporters, partial reconstitution of TP activity, even in a single organ will result in systemic clearance of excess nucleosides and clinical stabilisation or improvement in patients. While many patients affected by MNGIE now are offered hematopoietic stem cell transplantation (HSCT) or liver transplantation (LT), these procedures require immunosuppression and are limited by donor availability. Moreover, they are highly invasive interventions, with substantial peri– and post-procedure morbidity and mortality in this often severely affected patient group (*19–21*). Nevertheless, all reported patients who successfully underwent and survived LT or HSCT showed a sustained reduction of plasma dThd/dUrd to close-to-normal levels (*19, 20, 22–24*), and of those patients who lived 2 years post-HSCT, all showed a stabilization of many of the disease symptoms, with improvement of body mass index (BMI) and nerve conduction tests (*19, 23*).

Hepatotropic AAV-mediated gene therapy has been shown to ameliorate disease phenotypes in various MDS animal models, including in DGUOK, MPV17, TK2 or ANT deficiencies and in MNGIE (*25–30*). Despite this potential, there are currently no clinically approved AAV treatments for MDS. Barriers to approval for AAV gene therapies include lack of efficacy due to pre-existing antibodies, the development of neutralizing antibodies, AAV-gene silencing, reported deaths from acute liver failure and insertional oncogenesis (*31–35*). Preclinical AAV gene therapy studies for MNGIE in mouse models (*26, 36, 37*) or *ex situ* perfused liver explants (*38*), and case reports of successful liver transplants in patients (*22, 23*), demonstrate that nucleoside clearance can be achieved and disease phenotypes improved if hepatic expression is restored.

Intravenously (IV) administered lipid nanoparticles (LNPs) efficiently deliver mRNA-encoded proteins to the liver (*39, 40*) and transiently restore metabolic function in mouse models of monogenic metabolic disorders (*41–43*) and in a clinical trial of patients with propionic acidemia (*44*). However, mRNA-LNP protein replacement therapies require repeat administration, which for IV infusions can take many hours and need to be given under professional supervision. This places substantial burden on both medical professionals and patients, highlighting the need for better, safer and more accessible mRNA-LNP administration routes.

Here we demonstrate that *hTYMP*-mRNA-LNP-mediated reconstitution of TP expression in a mouse model of MNGIE completely restores systemic dThd/dUrd nucleoside balance. Through extensive biodistribution and pharmacodynamic studies, we find that IV and subcutaneous (SC) delivery of *hTYMP*-mRNA-LNPs induce hepatic TP expression and nucleoside clearance within two hours post-administration, with expression and therapeutic effects lasting up to 21 days. To improve SC delivery efficiency, we developed a novel approach by co-delivering mRNA-encoded human hyaluronidase. This demonstrates the potential of effective mRNA-based protein replacement therapy for primary mitochondrial disease, with proof-of-concept for biochemical efficacy after IV and SC administration of a clinically used LNP formulation. This approach provides a well-tolerated, accessible and directly translatable treatment strategy for patients affected by MNGIE, and can be extended to treat other MDS such as DGUOK, MPV17 and TK2 deficiencies, opening new therapeutic avenues for these fatal, but in principle curable diseases.

## RESULTS

### Synthetic *hTYMP*-mRNA delivery induces TP expression in liver cells

To restore TP expression in MNGIE, we designed a codon-optimized and uridine-depleted mRNA template encoding human *TYMP* (*hTYMP*) with UTRs from stable and highly translated mRNAs (*45, 46*) (**Fig. 1A**). A previously designed *eGFP*-mRNA (*47*) was used as control. *hTYMP*– and *eGFP*-mRNA were synthesized from cell-free DNA templates and co-transcriptional capped and tailed. To determine if synthetic *hTYMP-*mRNA induced robust TP protein expression, we transfected HuH-7 liver cancer cells. *hTYMP-*mRNA was strongly translated after 24 hours, with little impact on the expression of other proteins when compared to *eGFP*-mRNA transduced HuH-7 cells (**Fig. 1B, C; Table S1**).

**Figure 1.**
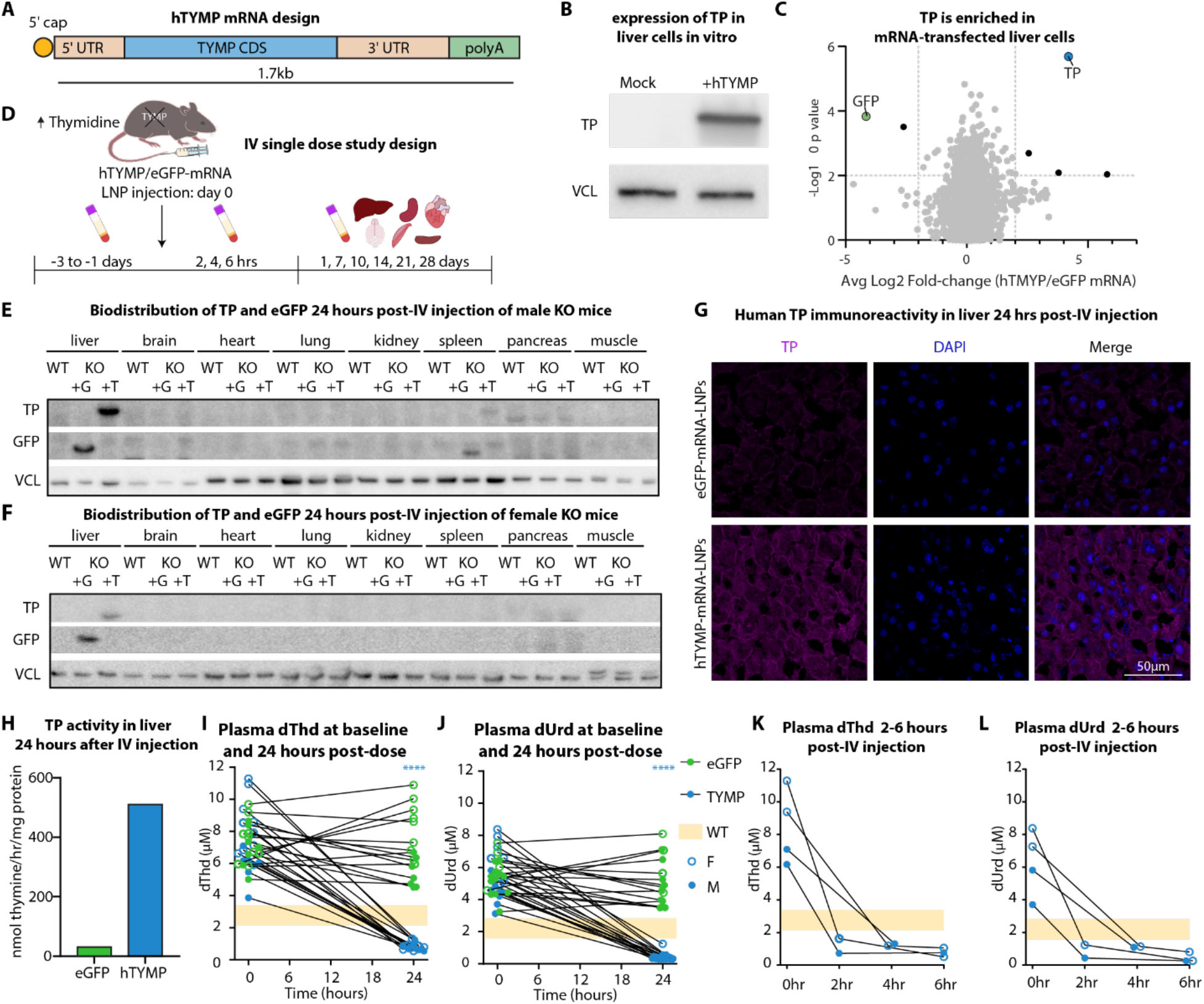
Efficacy of IV *hTYMP*-mRNA-LNP administration in MNGIE mice. **(A)** Schematic of synthetic *hTYMP*-mRNA produced by *in vitro* transcription. **(B)** TP and GFP Western blot in HuH-7 cells after mock or *hTYMP*-mRNA transfection. **(C)** TP and eGFP peptides identified by mass spectrometry (LC/MS). Dotted lines indicate FDR 0.01 (y-axis) and Log_2_ fold change ratio of 2 (x-axis). **(D)** Study design for single dose IV delivery of mRNA-LNPs to MNGIE mice. Plasma was collected prior to dosing (baseline) and three times after dosing, up to 28 days. Tissues were collected at weekly intervals with additional serum samples at baseline (–3 to –1 days), 2,4,6 hours and 10 days. **(E, F)** Western blots with TP and eGFP in the liver and spleen of male **(E)** and the liver of female **(F)** MNGIE (KO) or wild-type (WT) mice 24 hours after IV administration of 0.5mg/kg *eGFP*-mRNA-LNPs (+G) or *hTYMP*-mRNA-LNPs (+T). **(G)** TP immunostaining and DAPI nuclear stain in liver tissue 24 hours after IV injection. **(H)** TP activity in liver tissue from *hTYMP*– or *eGFP*-mRNA-LNP treated MNGIE mice, n = 1 per column. **(I, J)** dThd **(I)** and dUrd **(J)** levels in *hTYMP-* (blue) or *eGFP*– (green) mRNA-LNP treated female (open circles) and male (closed circles) MNGIE mice at baseline and 24 hours after injection. Two-way repeated measures ANOVA with uncorrected Fisher’s least significant difference; **** = p ≤ 0.0001, n = 16 per treatment. **(K, L)** dThd **(K)** and dUrd **(L)** levels in MNGIE mice at baseline, 2, 4 and 6 hours after injection of 0.5mg/kg *hTYMP*-mRNA-LNPs. **(I-L)** Each line indicates repeated sampling from one mouse; orange shading represents the range of nucleoside levels in wild type mice (WT).

### *hTYMP-*mRNA-LNP-induced hepatic TP expression restores systemic nucleoside balance in MNGIE mice

To evaluate mRNA-mediated TP protein replacement therapy as a treatment for MNGIE we used an established mouse model of MNGIE. The MNGIE mouse model is null for both *Tymp* and uridine phosphorylase 1 (*Upp1*) as *Upp1* also catabolizes thymidine (dThd); deletion of Tymp alone is therefore insufficient to produce the nucleoside accumulation that characterizes the human disease (*48, 49*). MNGIE mice are homozygous viable, but have elevated levels of plasma dThd and dUrd compared to wild-type mice (*9*), providing an ideal model to evaluate the biochemical efficacy of mRNA-LNP-based protein replacement therapy.

We first administered mRNA-LNPs encoding either *hTYMP* or *eGFP* intravenously at 0.5mg/kg (*42, 47*) and monitored protein, mRNA, and plasma nucleoside levels up to several weeks after injection. TP was strongly expressed in the livers of male and female *hTYMP*-mRNA-LNP treated mice and to a lower extent in the spleen of male mice, but not in other organs (**Fig. 1E, F**). A similar biodistribution was observed for eGFP protein, consistent with previous reports (**Fig. 1E, F**) (*39, 47*). TP and eGFP expression in the liver of treated animals were confirmed by immunofluorescence revealing widespread transfection of most hepatocytes, with TP primarily localized in the cytoplasm in concordance with TP’s native intracellular localization (**Fig. 1G, Fig. S1A**) (*50, 51*).

To evaluate whether *hTYMP*-mRNA-LNP administration resulted in biochemical rescue of the TP deficiency, we measured TP enzyme activity in homogenized liver tissue (*52*) from *hTYMP*-mRNA-LNP or *eGFP*-mRNA-LNP injected MNGIE mice. High TP activity was detected in the liver of both male and female mice 24 hours post-treatment **(Fig. 1H, Fig. S1B)**. Lower activity was observed in spleen, with no or barely detectable activity in tissues of *eGFP*-treated MNGIE mice (**Fig. 1H, Fig. S1C**). Crucially, high liver TP activity 24 hours after *hTYMP*-mRNA-LNP injection significantly reduced circulating dThd and dUrd levels to concentrations well below the levels measured in wild type mice (**Fig. 1I, J; Table S2**). Nucleoside depletion was specific to TP activity, as *eGFP*-mRNA-LNP administration did not affect plasma nucleosides compared to baseline levels (**Fig. 1I, J**). To determine early dynamics of nucleoside clearance, we measured plasma nucleoside levels at two, four and six hours, and observed a significant reduction to below wild-type levels within two hours after *hTYMP*-mRNA-LNP injection (**Fig. 1K, L**). Together, the strong TP expression and early nucleoside clearance data provide proof-of-concept for systemic *hTYMP*-mRNA-LNP-based protein replacement therapy in a model for MNGIE, demonstrating liver-specific expression, and rapid biochemical efficacy on a clinically relevant biomarker.

### Long-term efficacy of mRNA-derived TP expression

We next assessed long-term efficacy and pharmacodynamics of IV *hTYMP*-mRNA-LNP administration by measuring *hTYMP* mRNA and TP protein expression in the liver, and plasma dThd/dUrd levels up to four weeks post-injection. *hTYMP* mRNA was only detectable in liver tissue in the first 24 hours after injection, and became undetectable afterwards (**Fig. S2A**). However, robust TP protein expression (**Fig. S2B**) and activity (**Fig. S2C**) were maintained in the livers of male and female mice at seven days after injection, with further low-level expression at seven days in spleen. Hepatic TP expression declined in the second week, but remained clearly detectable up to 14 days, and continued at lower levels in both sexes up to 21 days after injection (**Fig. 2A**), before becoming undetectable at 28 days (**Fig. S2D**). TP expression was strongest at 24 hours and by 21 days had decreased to 5.7% ± 6.6 (mean ± SD) in males and 2.3% ± 1.1 (mean ± SD) in females of this peak expression (**Fig. 2B**). We calculated *in vivo* apparent TP decay half-time in mouse liver to be between 4.46 days (males) and 6.18 days (females) (**Fig. 2B**). Mean systemic dThd/dUrd nucleoside levels remained below or within wild type levels until at least 14 days postinjection (**Fig. 2C, 2D; Table S2**), and only started to increase after two weeks, to reach baseline levels again by 21 days. Together, the strong TP expression and early pharmacodynamic biomarker data demonstrate that a single dose of *hTYMP*-mRNA-LNPs allows for sustained (> 2 weeks) hepatic TP expression and restoration of nucleoside balance in MNGIE mice, providing the preclinical data required to inform future repeat dosing studies in mice and patients.

**Figure 2.**
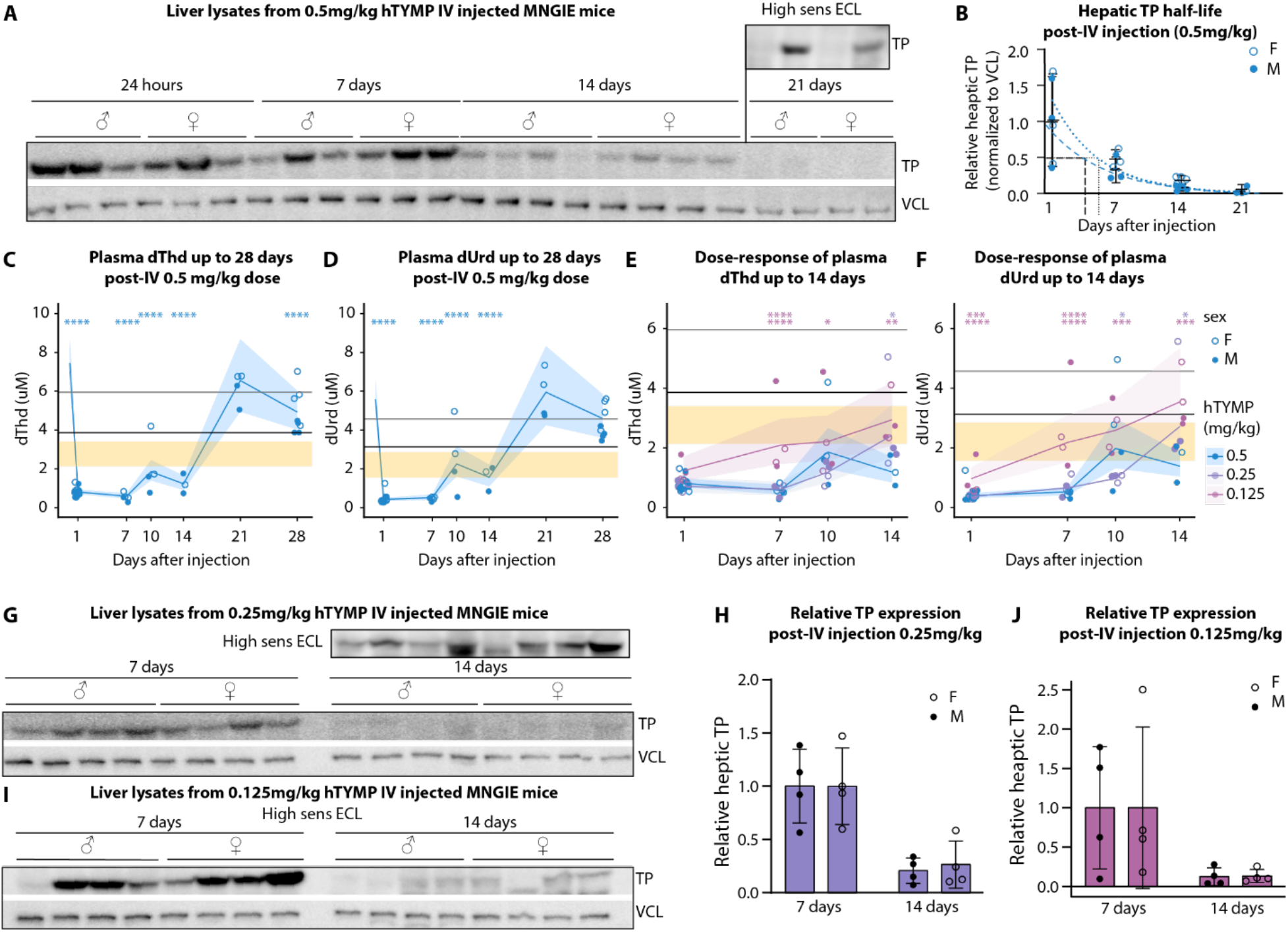
Prolonged and dose-dependent efficacy of *hTYMP*-mRNA-LNPs. **(A)** Western blot of hepatic TP expression up to 21 days after IV 0.5mg/kg *hTYMP*-mRNA-LNP injection in male and female MNGIE mice. **(B)** Normalized hepatic TP expression from levels detected by Western blot in **(A)**. Estimated protein half-life in males (blue dashed line) and females (blue dotted line). **(C-F)** Estimated marginal mean plasma dThd **(C, E)** and dUrd **(D, F)** levels in MNGIE mice up to 14 **(E, F)** or 28 **(C, D)** days after IV administration of indicated doses of *hTYMP*-mRNA-LNPs. (**G-J**) Western blot **(G, I)** and quantification **(H, J)** of TP expression in liver at 7 and 14 days after 0.25mg/kg (**G, H**) or 0.125mg/kg **(I, J)** *hTYMP*-mRNA-LNPs IV injection. **(B-F, H, J)** Each dot represents one mouse. Graphs (**C-F**) show estimated marginal means (lines) and 95% confidence intervals (shading) from generalized linear mixed-effects models with mouse as a random effect and pairwise contrasts of estimated marginal means, corrected for multiple comparisons via the Holm method. * = p ≤ 0.05, ** = p ≤ 0.01, *** = p ≤ 0.001, **** = p ≤ 0.0001. Grey (female) and black (male) lines indicate the baseline lower limit of nucleoside concentrations in MNGIE mice; orange shading indicates the range of nucleoside concentrations in wild type mice.

### hTYMP-mRNA-LNPs restore nucleoside balance in a dose-dependent manner

To determine the preclinical minimally effective dose range for nucleoside balance restoration, we conducted a two-week study of MNGIE mice treated with 0.25mg/kg or 0.125mg/kg *hTYMP*-mRNA-LNPs. Plasma dThd/dUrd levels (**Table S2**) rapidly reduced to below the levels detected in wild-type mice with both doses at 24 hours, similar to those observed with 0.5mg/kg mRNA-LNPs. Mean dThd and dUrd levels remained at or below wild-type levels for up to 10 days after injection, but were consistently higher in mice injected with the lowest dose (0.125mg/kg; compared to 0.25mg/kg or 0.5mg/kg) (**Fig. 2E, F**), indicating a clear dose-responsive effect of *hTYMP*-mRNA-LNP treatment. However, at 14 days, dThd/dUrd levels in most mice treated with 0.25mg/kg still remained within wild type levels (**Fig. 2E, F**), potentially indicating a preclinical therapeutic dose range between 0.25mg/kg and 0.5mg/kg, with 0.25mg/kg being the lowest tested dose still providing significant biochemical efficacy after 2 weeks (0.5mg/kg vs 0.25mg/kg p = 0.0107 (dThd), p = 0.0169 (dUrd); 0.5mg/kg vs 0.125mg/kg p = 0.0011 (dThd), p = 0.0007 (dUrd); 0.25mg/kg vs 0.125mg/kg p = 0.443 (dThd), p = 0.367 (dUrd)). In line with these dose-dependent dThd/dUrd reductions, both dose levels sustained hepatic TP expression up to 14 days post-injection, but more robustly so at 0.25mg/kg (**Fig. 2G-J**), together informing future clinical dose-finding studies.

### *hTYMP*-mRNA-LNPs are well tolerated with transient side effects

mRNA-LNP-based protein replacement therapies are generally assumed to be safer alternatives to virus-mediated gene therapy (*53*) but must be compatible with repeated dosing. We therefore evaluated systemic toxicity after IV administration of *hTYMP*– and *eGFP*-mRNA-LNPs. Both MNGIE and wild-type groups treated with either mRNA-LNP exhibited a small (<5%) transient decrease in body weight after 24 hours compared to baseline and to sham-injection (**Fig. 3A; Table S3**). This remained within local welfare guidelines, was not dose-dependent, started to recover by 48 hours post-injection, and had returned to baseline by seven days (**Fig. 3B-D**).

**Figure 3.**
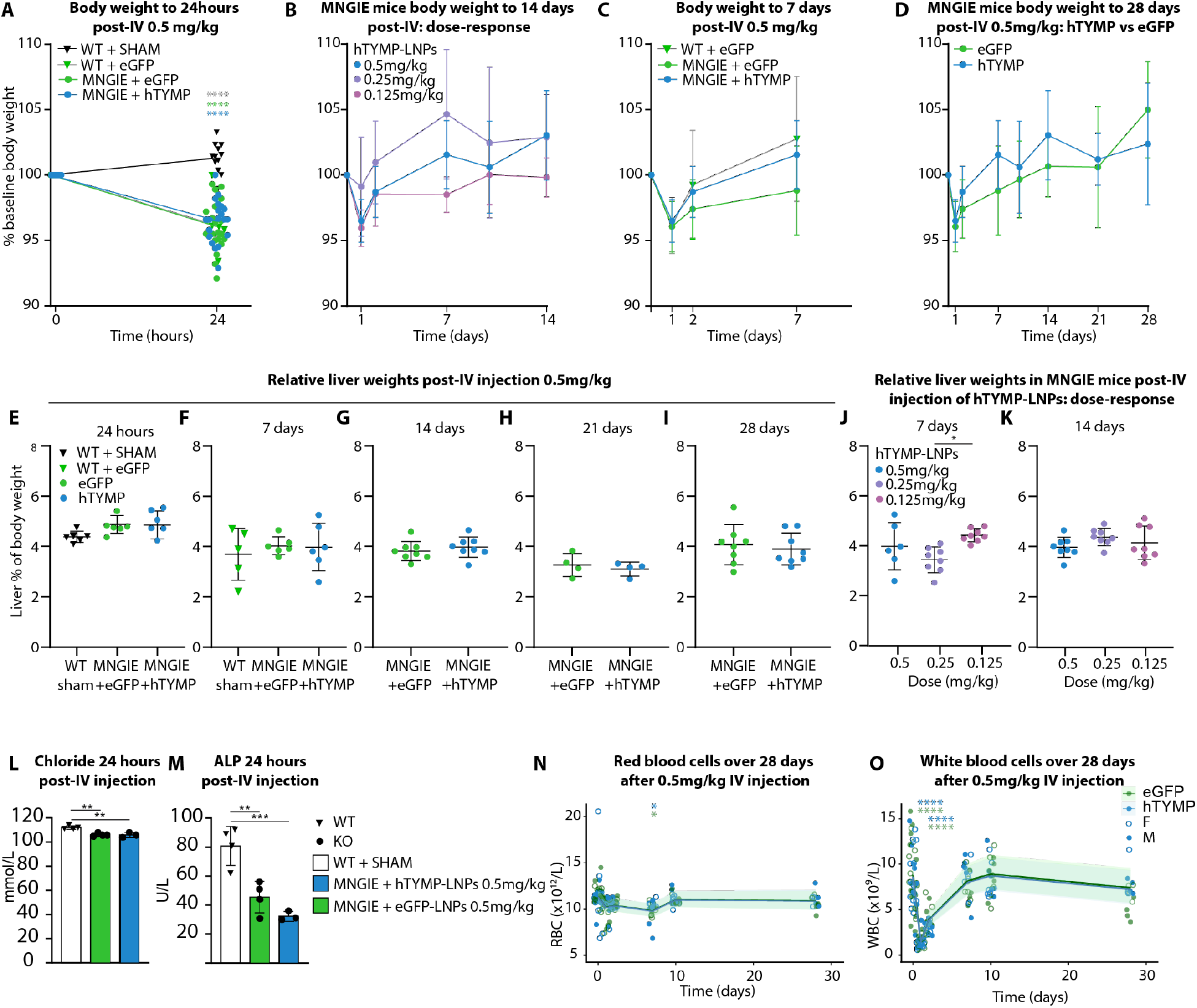
IV administration of mRNA-LNPs is well tolerated. **(A)** Mean body weight at 24 hours after IV administration of mRNA-LNPs compared to sham-injected mice. WT sham n = 12, WT + *eGFP*-mRNA-LNPs n = 6, MNGIE + *hTYMP*-mRNA-LNPs n = 20, MNGIE + *eGFP*-mRNA-LNPs n = 20. **(B)** Mean body weight after IV delivery of 0.125mg/kg (pink), 0.25mg/kg (purple) or 0.5mg/kg (blue) *hTYMP*-mRNA-LNPs. **(C, D)** Mean body weight up to 7 **(C)** or 28 **(D)** days of *hTYMP*-(blue) or *eGFP*-(green) mRNA-LNP treated wild-type (triangles) or MNGIE (circles) mice. **(E-I)** Relative liver weights up to 28 days after IV sham injection **(E)** or 0.5mg/kg *eGFP-* (green) or *hTYMP*-(blue) mRNA-LNPs (E-I). **(J, K)** Relative liver weights at three doses of *hTYMP*-mRNA-LNPs after 7 **(J)** and 14 **(K)** days. **(L, M)** Chloride **(L)** and ALP **(M)** levels 24 hours post-injection. **(N, O)** Peripheral red blood cell **(N)** and white blood cell **(O)** counts up to 28 days after IV *eGFP*-(green) or *hTYMP*-(blue) mRNA-LNP delivery. Data in **(B-K)** are represented as mean ± SD. **(A, E-Q)** Dots represent individual mice; see **Table S3** for n-numbers in (**B, D**). One-way ANOVA with Dunnett’s multiple comparisons **(A)**; One-way ANOVA with Tukey’s multiple comparisons test **(E, F, J-M)**; Unpaired t-test **(G-I)**; Estimated marginal means (lines) with 95% confidence intervals (shading) determined by generalized linear mixed-effects models with mouse as a random effect and pairwise contrasts, corrected for multiple comparisons via the Holm method **(N, O)**. * = p ≤ 0.05, ** = p ≤ 0.01, *** = p ≤ 0.001, **** = p ≤ 0.0001. WT = wild type mice.

Relative liver weight, as an indicator of hepatic injury (*54*), did not differ between sham– or mRNA-LNP injected wild type and MNGIE mice at all time-points tested (**Fig. 3E-K**). Most analytes from a broad biochemical serum panel showed no significant differences between mRNA-LNP treated MNGIE mice and sham-injected wild type mice from 24 hours up to 28 days after IV injection, apart from decreased chloride and ALP levels in mRNA-LNP-treated mice (**Fig 3L, M, Fig. S3A-C; Table S4**), which had recovered at 7 days (**Fig. S3B**), providing further evidence for limited organ impact of systemic mRNA-LNP administration (*44, 55*), without signs of TP-related toxicity.

Small decreases in red blood cell indices occurred up to seven days, then recovered and remained stable up to 28 days (**Fig. 3N, Fig. S4A-F; Table S5**). White blood cell counts, including lymphocytes and neutrophils, as well as platelets, decreased significantly as soon as 24h after IV injection, in both *hTYMP*-mRNA-LNP and *eGFP*-mRNA-LNP treated mice (**Fig. 3O, Fig S4G-L; Table S5**). This leukopenia was not significantly dose-dependent and was also present in wild-type mice injected with eGFP-mRNA-LNPs (**Fig. S5**), indicating a payload– and genotype-independent mRNA-LNP effect. White blood cell counts started to recover rapidly, within 48 hours after injection, and returned to baseline within seven days (**Fig. 3O**). Together, these preclinical safety data demonstrate transient reductions in total body weight and circulating white blood cells, independent of TP expression but related to the LNP platform, offering scope for further formulation optimization. However, IV administration of *hTYMP*-mRNA-LNPs in MNGIE mice is otherwise well tolerated, without evidence for transgene-related toxicity.

### Subcutaneous administration of *hTYMP*-mRNA-LNPs rescues systemic nucleoside levels

Having established preclinical safety and efficacy of systemic *hTYMP*-mRNA-LNP administration for the treatment of MNGIE, we next sought to develop a less invasive and simpler method of drug delivery than IV administration. This is particularly important for therapies with transient effects like mRNA-LNPs that require repeat-administration. SC delivery is only recently (*56*) being explored as an administration route for mRNA-LNP-mediated protein replacement therapies. SC mRNA-LNP injections could be self-administered, outside of a clinic setting, reducing the burden on patients, carers, and healthcare services. We therefore investigated if SC *hTYMP*-mRNA-LNP administration could induce sufficient TP expression in skin, liver or other organs to effectively reduce systemic nucleoside levels (**Fig. 4A**). SC *hTYMP*– or *eGFP*-mRNA-LNPs injection induced variable but dose-dependent hepatic TP protein expression. 0.5mg/kg *hTYMP*-mRNA-LNPs only induced expression in skin tissue near the injection site, but higher mRNA-LNP doses, at 1mg/kg and 2mg/kg, resulted in reproducible hepatic GFP or TP expression (**Fig. S6A, Fig. 4B, C**). We noticed a tendency towards higher TP expression in female mice, possibly reflecting sex-dependent differences in skin composition (*57, 58*). Hepatic TP expression remained detectable until at least seven days post-injection (**Fig. 4D**), but with variable efficiency and at lower levels than after IV administration. Systemic dThd/dUrd nucleoside levels were reduced to wild-type levels within 24h after SC injection for most treated mice (**Fig. 4E, F; Table S2**). Although strict interpretation of nucleoside levels was confounded by large variation in the measurement of baseline dThd levels, these results demonstrate the feasibility of SC mRNA-LNP administration.

**Figure 4.**
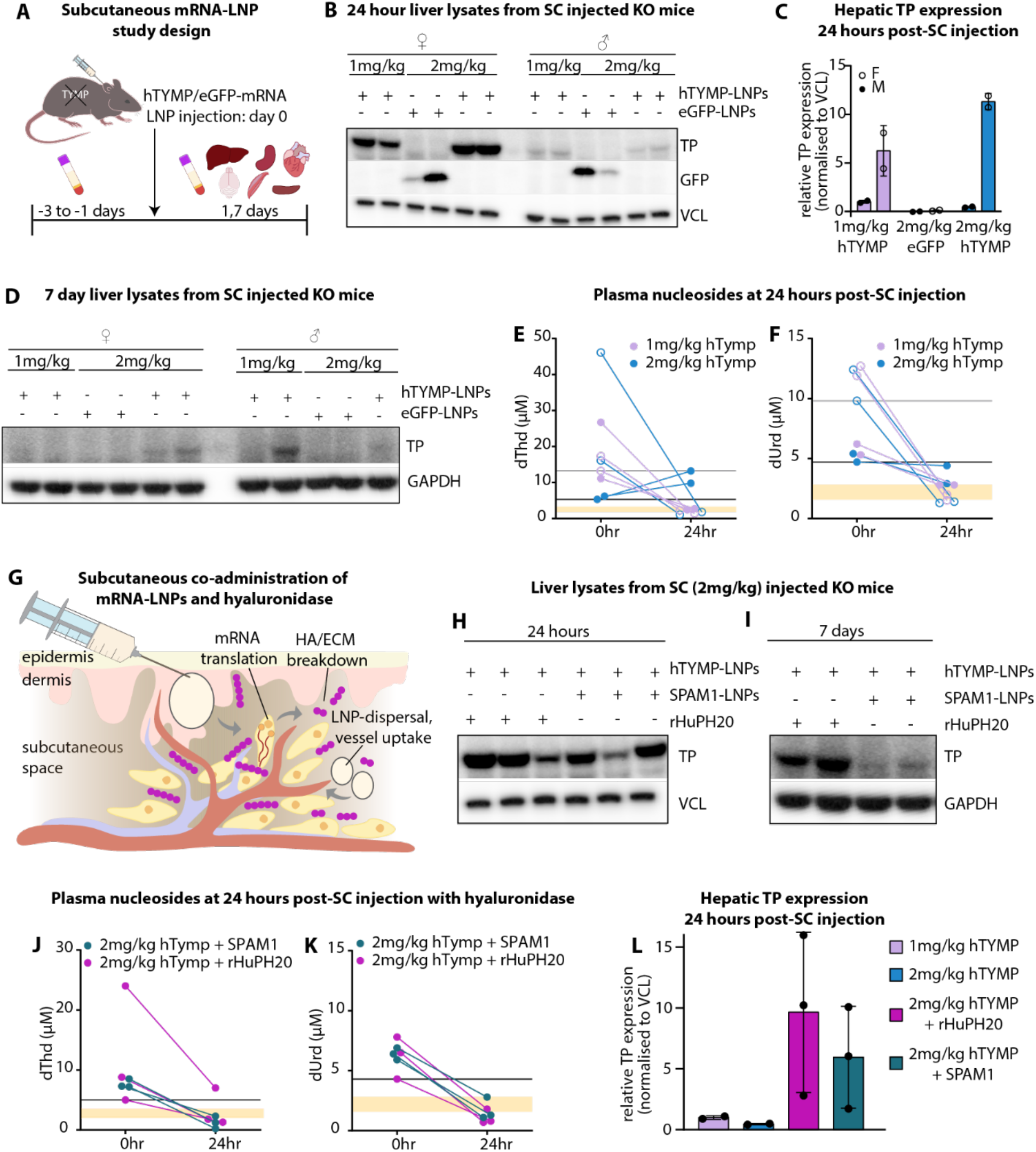
SC administration of *hTYMP*-mRNA-LNPs enables hepatic expression and reduces nucleoside levels in MNGIE mice. **(A)** Study design for SC delivery of mRNA-LNPs in MNGIE mice. Plasma was collected prior to dosing and up to three times over 7 days after injection. Tissues were collected 24 hours and 7 days after injection. **(B-D)** Western blot **(B, D)** and quantification **(C)** of hepatic TP expression 24 hours **(B,C)** or 7 days **(D,L)** after SC *hTYMP*-mRNA-LNP delivery in MNGIE mice. **(E,F)** Plasma dThd **(E)** and dUrd **(F)** levels 24 hours after injection. The lower limits of nucleosides prior to treatment are indicated by horizontal grey (female) and black (male) lines. Orange shading indicates nucleoside levels in wild-type mice. **(G)** Hyaluronidase breaks down hyaluronic acid (HA) in the ECM of the subcutaneous space facilitating LNP uptake into surrounding vessels**. (H)** Western blot of liver tissue 24 hours and seven days **(I)** after co-administration of SC *hTYMP*-mRNA-LNPs with hyaluronidase. **(J, K)** Plasma dThd (**J**) and dUrd **(K)** 24 hours after SC co-administration of 2mg/kg *hTYMP*-mRNA-LNPs with 2mg/kg *SPAM1*-mRNA-LNPs (teal) or rHuPH20 (magenta). Lower limits of nucleosides prior to treatment are indicated by a horizontal black line, orange shading indicates nucleoside levels from wild-type mice. Dots represent individual mice.

### Hyaluronidase enhances subcutaneous mRNA-LNP delivery

Because SC efficacy of mRNA-LNPs was lower than upon IV injection, we next investigated whether concomitant delivery of hyaluronidase, to transiently modify the extracellular matrix at the injection site, could enhance LNP uptake in the surrounding vasculature (**Fig. 4G**) (*56, 59*). We co-administered 2mg/kg *hTYMP*-mRNA-LNPs with 2mg/kg *SPAM1*-mRNA-LNPs encoding human hyaluronidase or with recombinant engineered human hyaluronidase (rHuPH20). Male mice were used, since these showed lowest TP expression after standard SC administration (**Fig. 4C**). Co-administration of 2mg/kg *hTYMP*-mRNA-LNPs with either *SPAM1*-mRNA-LNPs or rHuPH20 increased hepatic TP protein expression 24 hours after injection (**Fig. 4H**). Hepatic TP expression was sustained until at least seven days after SC administration, with strongest expression observed in those co-treated with rHuPH20 (**Fig. 4I**). Plasma dThd/dUrd were rapidly reduced to or below wild-type levels for all mice treated with 2mg/kg *hTYMP*-mRNA-LNPs and *SPAM1*-mRNA-LNPs and in two of three mice treated with 2mg/kg *hTYMP*-mRNA-LNPs and rHuPH20, although variability in baseline nucleoside levels in these SC experiments prevented us from drawing clear conclusions about reproducibility of the efficacy (**Fig. 4J-L; Table S2**).

Co-administration of rHuPH20 or *SPAM1*-mRNA-LNPs was associated with short-term toxicity, illustrated by lower total body weight and relative liver weight after 24 hours compared to animals and livers that only received *eGFP*– or *hTYMP*-mRNA-LNPs (**Fig. S6B, C**). In addition, at 24 hours, livers from animals treated with *SPAM1*-mRNA-LNPs were paler and softer to dissect **(Fig. S6D).** While total body weight had fully recovered after seven days (**Fig. S6B**), liver weight remained lower in hyaluronidase-treated animals (**Fig. S6E**). Serum analytes (**Fig. S6F; Table S4**) and whole blood composition (**Fig. S7; Table S5**) did not show additional changes attributable to hyaluronidase co-delivery over 7 days after injection. Together, these data demonstrate the feasibility of hyaluronidase co-administration to enhance efficacy of SC mRNA-LNP-mediated transgene delivery, with short-term tolerability considerations that will require future optimization for repeat administration.

## DISCUSSION

MNGIE is an ultra-rare mitochondrial disease caused by autosomal recessive loss-of-function mutations in *TYMP.* MNGIE is rapidly progressive and fatal, but in principle curable. Because dThd and dUrd, the toxic metabolites that drive disease progression, readily equilibrate across the body, successful treatment of this multisystem condition can be achieved by correcting TP expression in a single organ only. This creates a unique opportunity for current organ-specific non-viral protein replacement strategies.

Using a biochemically relevant MNGIE mouse model, we demonstrate that a single dose of IV *hTYMP*-mRNA-LNPs and the resulting hepatic TP expression, reduces nucleoside levels to below levels detected in wild type mice, for a sustained period up to 14 days after administration. This is in line with previous human findings, where modest TP protein expression, either in heterozygous family members, in late-onset MNGIE patients with partial TP activity, or upon organ transplantation, can maintain systemic nucleoside balance and prevent or delay disease progression (*14, 15, 19, 22*). We define apparent hepatic TP decay half-time and the preclinical minimally effective IV dose range for *hTYMP*-mRNA-LNPs, which together will inform future repeat dosing studies in mice and patients. Furthermore, we demonstrate the first biochemical rescue in a mitochondrial disease model after SC administration of an mRNA therapy and achieve enhanced hepatic TP expression by co-administration of hyaluronidase protein or mRNA.

SC delivery is a preferred administration route by patients and care givers when compared to IV delivery (*60–62*). SC injection allows self-administration, improves patient quality of life, and multiple sites can be used to facilitate repeated delivery, which is crucial for transient protein-replacement approaches like mRNA-LNPs (*63*). We observed differences in hepatic TP expression and nucleoside clearance between male and female mice after SC injection, possibly related to sex-specific dermal and hypodermal layer thickness (*57, 58*). Although sex differences in dermal thickness at common SC injection sites in humans are less extreme and overcome by needle length, sex-specific responses should be documented as these differences in dermal composition may still influence pharmacokinetics of subcutaneously delivered drugs (*64, 65*).

Subcutaneous co-administration of hyaluronidase, to transiently break down the extracellular matrix component hyaluronic acid, can improve drug absorption and increase injection volumes in patients with primary immune deficiencies and some cancers (*59, 66–68*). In patients with type II diabetes, SC co-delivery of rHuPH20 permits a reduced dose of the treatment-specific drug (*69*). Co-administration of rHuPH20 also reduced pro-inflammatory cytokine levels in mice repeatedly dosed SC with mRNA-LNPs, which is important for conditions like MNGIE where repeat dosing is required (*56*). We find that co-administration of hyaluronidase, either as rHuPH20 or as *SPAM1*-mRNA-LNPs, together with *hTYMP*-mRNA-LNPs enhances hepatic TP expression and nucleoside reduction in MNGIE mice. *SPAM1*-mRNA-LNPs provide an attractive alternative to rHuPH20, being manufactured on the same platform and administered in the same formulation as the therapeutic mRNA-LNP, allowing transient and tunable expression. Although hyaluronidase administration led to a decrease in relative liver weights, possibly due to hyaluronidase activity in the liver, *SPAM1*-mRNA-LNPs could be engineered to prevent hepatic SPAM1 expression but remain active at the injection site, for example by inclusion of a miRNA binding site in *SPAM1*-mRNA. Dual mRNAs have been repeatedly delivered safely in a Phase1/2 clinical trial of propionic acidemia (*44*) further supporting clinical translation of our approach.

We estimated the apparent decay half-life of human TP in the livers of MNGIE mice to be 4-6 days, reflecting optimal codon adaptation indexing during mRNA design, continuous protein production until mRNA degradation and the tendency for protein half-lives of abundant proteins to increase in higher order organisms (*46, 70*). The half-life of human TP in plasma after IV delivery in rats is seven hours, and of endogenous TP in mouse liver is 3.1 days (*71, 72*). However, low protein expression in other organs and the lack of *hTYMP*-mRNA detected in liver after 24 hours, indicate that prolonged hepatic TP expression after *hTYMP*-mRNA-LNP administration is likely due to the TP protein stability rather than the persistence of *hTYMP*-mRNA.

We also conducted extensive safety analysis and found no evident organ-specific toxicity related to the *hTYMP*-mRNA-LNP payload and transgene expression. In contrast, transient decrease in body weight and hematologic effects were found, likely attributable to the LNP platform. There are limited preclinical reports of the acute effect of LNPs on circulating immune cells (*55, 73, 74*), and it will be important to assess broader relevance of the acute decrease in circulating white blood cells we observed after mRNA-LNP administration. The transient post-treatment leuko– and thrombocytopenia were seen with all doses, transgenes, genotypes and delivery routes, indicating a response to the carrier LNPs. These hematological side-effects could be due to immune-cell translocation to the liver, as seen after intradermal injection of Acuitas-LNPs in preclinical models (*75*). Thrombocytopenia has also been observed in clinical trials with LNP-based medicines (*76, 77*) leading to LNP reformulation (*76*). We chose SM102 as an LNP-formulation approved for use in humans which facilitates strong protein expression after IV delivery, mRNA bioavailability after SC delivery, and reduced activation of the innate immune system compared to other clinically approved ionizable lipids such as ALC-0-315 (*40, 47, 78*). Transient peripheral leukopenia might be avoided in future studies by changing the ionizable lipid. We did not observe other organ-specific toxicity, such as necrosis of liver cells and elevated transaminases (ALT, AST) previously observed after treatment of rats with empty MC3-LNPs (*79*). While robust and sustained protein expression was enabled by SM102-LNPs after both IV and SC delivery, much remains to be done in the field of LNP development to avoid these transient hematological side effects.

Unlike viral vector mediated gene therapy, mRNA therapy can be withdrawn if toxicities develop, or the dose increased if initial dosing is insufficient. Compared to recombinant proteins used in enzyme replacement therapies, mRNA therapies are protected from degradation after administration by their LNP shells and can act intracellularly in targeted tissues. Needing access only to the cytoplasm, mRNAs can change cell responses rapidly, as demonstrated by the sharp drop in nucleoside levels we observed two hours after delivery in the MNGIE mouse model. Clinical translation will depend on confirmation of comparable pharmacodynamics and safety profiles in humans, and on further optimization of the LNP-platform for repeat administration. In particular, LNP modification may prevent transient leukopenia, and liver detargeting of *SPAM1*-mRNA may enhance SC delivery while preventing hepatic hyaluronidase expression. The sustained biochemical correction, for at least 14 days, supports the feasibility of repeat administration, including through SC self-administration at home, with the optimal dosing interval to be established in future repeat-dose preclinical and clinical studies. Together, these findings support the development of mRNA-LNP-mediated TP replacement as a repeatable approach to maintaining systemic nucleoside homeostasis in MNGIE and other MDS, without the risks associated with viral gene delivery or the need for lifelong immunosuppressants.

## MATERIALS AND METHODS

### Study Design

The objective of this study was to investigate if delivery of LNPs encapsulating mRNA-encoded *hTYMP* was an effective approach for TP protein replacement therapy in a mouse model of MNGIE. Animal studies were designed with equal numbers of age and sex matched MNGIE mice for mRNA-LNP injections, except for the combination hyaluronidase study which was conducted in male mice. Control *eGFP*-mRNA-LNPs and test-mRNA-LNPs (*hTYMP*, *SPAM1*) were injected on the same day, and sample collections from each condition were performed on the same day in a randomised order. Plasma was analysed to monitor systemic nucleoside levels. Tissues were measured for mRNA-derived TP expression and activity. One plasma sample was unable to be collected from a male mouse 6 hours after IV injection of 0.5mg/kg *hTYMP*-mRNA-LNPs; all other samples and analyses from the study are included in the manuscript and supplementary tables. For analysis of plasma nucleosides at least 3 samples were analysed at each timepoint.

### mRNA design and production

mRNA production was performed by the BASE facility at The University of Queensland. In brief, mRNA was produced from a linearised plasmid template using *in vitro* transcription and co-transcriptional capping incorporating N1-methylpseudouridine. For encapsulation aqueous mRNA was mixed with lipids diluted in ethanol on a NanoAssemblr Ignite followed by dialysis with phosphate-buffered saline (PBS). LNPs were composed of SM-102, cholesterol, DSPC and DMG-PEG2000 in a ratio of 50:38.5:10:1.5. Size, PDI and Zeta potential were measured on a Malvern Zetasizer. Mean particle sizes were 69.73nm (*eGFP*-mRNA-LNPs), 73.69nm (*hTYMP*-mRNA-LNPs) and 79.93nm (*SPAM1*-mRNA-LNPs).

### *in vitro* protein expression and mass spectrometry

HuH-7 cells were grown in 6 well plates with complete DMEM (119065092 ThermoFisher, 10% fetal bovine serum and penicillin-streptomycin) and transfected with 500ng mRNA using MessengerMAX 24 hours prior to freezing cell pellets. Cell pellets were resuspended in RIPA buffer plus HALT protease inhibitors for western blots or processed by Q-MAP staff at The University of Queensland for mass spectrometry (MS). MS samples were prepared via S-Trap Micro Spin Column Digestion performed with SDS lysis buffer and DTT to a final concentration of 20mM with subsequent trypsin digestion. The S-trap was loosely capped and incubated at 37°C overnight. Peptides were eluted three times with 40 µL of 5%/50%/75% ACN in 0.1% FA, respectively, and eluent was dried in a speedy vac. Prior to liquid chromatography-mass spectrometry samples were redissolved in 20 µl of buffer A. LC-MS/MS was performed using a UHPLC system (Thermo Fisher Scientific) coupled to an Exploris 480 mass spectrometer with a FAIMS Pro interface using standard settings. Data analysis was performed using Spectronaut against the human proteome and eGFP sequence with a Q-value cutoff of 0.05 (https://www.uniprot.org/proteomes/UP000005640).

### MNGIE mouse model

Animals were housed in a Home Office-designated facility, according to the UK Home Office guidelines upon approval by the University of Cambridge Animal Welfare & Ethical Review Body (AWERB) and the UK Home Office (project license PP1740969 and PP8565009). Mice were kept in individually ventilated cages at 20-24°C, 45-65% humidity on a 12 h light/dark cycle with continual access to food and water. The mouse line used in this study was Tymp^-/-^/Upp1^-/-^, a double homozygous knockout of *Tymp* and *Upp1*, obtained from Vall d’Hebron Institut de Recerca (VHIR). C57BL/6J wild type mice were purchased from Charles River Laboratories.

### Protein detection by western blot

Tissue lysates were prepared by bead homogenisation of <30mg tissue at 5500 rpm for 15 seconds in PathScan CST buffer with Roche cOmplete protease inhibitors using a Precellys 24 Bead Mill Homogeniser. 10-30µg total protein as determined by BCA was run on NuPAGE Novex 4-12% BT Midi Gels and transferred using iBlot 2 PVDF transfer stacks for 7 minutes at 20V. Membranes were blocked and incubated with antibodies diluted in 5% BSA-0.1%TBSTovernight at 4°C. Membranes were washed in 0.1% TBST and incubated with secondary antibodies for one hour at room temperature, washed again and imaged on a GE ChemiDoc after incubation with Pierce ECL HRP substrate or Amersham ECL Prime Western Blotting Detection Reagent. The *in vivo* half-life of TP was calculated by fitting an exponential curve for protein degradation to the normalised TP protein signal and assuming no new protein synthesis after 24 hours. Antibody concentrations are listed in **Table S6**.

### Immunostaining and imaging

Snap-frozen livers were embedded in optimal cutting temperature compound, cryosectioned at 10µm and stored on Superfrost Plus slides at –80°C until immunostaining. Sections were fixed in 4% paraformaldehyde (PFA) for ten minutes or in a 1:1 acetone:methanol solution at –20°C for 7 minutes then air-dried at room temperature, followed by three PBS washes (+0.3% triton X-100 for PFA-fixed sections). Sections were blocked in 0.3% Triton-X100, 2% bovine serum albumin in PBS for two hours and incubated overnight at 4°C with primary antibodies diluted in block. The next day sections were washed as for post-fixation and quenched for lipofuscin autofluorescence for 45 seconds, incubated with secondary antibodies in PBS for two hours, followed by a 15-minute incubation with DAPI prior to three PBS washes and mounting with ProLong Glass Antifade Mountant. PFA fixation, washes and secondary antibody incubations were performed at room temperature. Images were captured by a Zeiss LSM 880 confocal scanning system. Antibody concentrations are listed in **Table S6**.

### TP activity assay

TP activity was measured as previously described (*52*) with minor modifications. Tissue homogenates containing 100 µg of protein were incubated in arsenate buffer in the presence of the TP substrate (10 mM thymidine) in 100 µL of reaction mix for one hour at 37°C, with subsequent inactivation of TP by protein precipitation by addition of 0.5 M perchloric acid. A parallel blank was processed in duplicate samples where thymidine was omitted during the incubation and added just after acidic inactivation of TP. The protein precipitate was pelleted by centrifugation and the thymine concentration was quantified in the supernatants by high performance liquid chromatography (HPLC) with UV detection. The result of the thymine concentration in the blank was subtracted from that quantified in the full reaction, and the TP enzyme activity results were expressed as nmoles of thymine formed/hour/mg protein.

### Plasma nucleoside detection

dThd and dUrd levels were measured by HPLC as previously described (*36*). Briefly, whole blood from the dorsal pedal vein was collected into EDTA-microvettes with 100µM TP inhibitor (6-Amino-5-bromo-1H-pyrimidine-2,4-dione) on ice and centrifuged at 3000g for five minutes at 4°C. Plasma was removed, diluted 1:5 in PBS and snap-frozen until analysis. At the time of analysis, proteins were removed from diluted plasma by ultrafiltration using 10 kDa filters (Amicon Ultra Filters, Millipore). The ultrafiltrates were injected in triplicates on an Acquity UPLC-Xevo TQ Mass Spectrometer (Waters, Milford, MA) and resolved in a gradient elution using a reverse-phase column (*36*). The analytes were detected by multiple reaction monitoring with positive electrospray using the following m/z transitions: 242.8>127.1 (dThd), 228.8>113.1 (dUrd) and 244.8>113.0 (rUrd). Calibration curves made with an aqueous multi-standard containing dThd, dUrd, rUrd (concentrations ranging 0.05 – 50 µM) were processed in parallel, and sample concentrations were obtained from interpolation of the peak areas corrected by the internal standard isotope-labeled ^13^C ^15^N –dThd, added to all standards and samples. Raw data for nucleoside levels are provided in **Table S2**.

### Hematology and clinical chemistry

20ul of whole blood was collected from mouse pedal veins and placed 1:25 in diluent prior to determination of hematological values via electrical impedance, colorimetric reactions and flow cytometry on an Element HT5Veterinary Hematology Analyzer. Serum samples for biochemistry were collected via cardiac puncture at termination. Blood was left to clot at room temperature for 20 minutes, centrifuged at 1300g for 15 minutes and the serum was flash frozen at –80°C until thawing for analysis via spectrophotometry and potentiometry on a Beckman Coulter AU480 Chemistry Analyzer. Raw data for serum biochemistries and hematological cell counts are in **Table S4** and **Table S5**.

### Statistical analysis

Blood samples collected longitudinally for nucleoside levels and hematology were analyzed using R via linear mixed-effects models (LMMs) or generalized linear mixed-effects models (GLMMs), with mouse and/or sample date included as random effects. Sample date was included as an additional random effect where its inclusion was supported by a likelihood ratio tests (LRT) comparing models with and without this term. LRTs assessed the significance of the dose × time interaction, comparing additive (dose + time) and interaction (dose × time) models. When the interaction was not significant, inference was based on the simpler additive model. Model distribution was selected for each outcome based on residual diagnostics using posterior predictive checks and the Akaike Information Criterion (AIC). P-values for pairwise comparisons were adjusted using Tukey or Holm corrections as appropriate.

Serum samples collected at 24 hours or seven days were analyzed via one-way ANOVA with Tukey’s multiple comparisons or Krukal-Wallis tests. Comparison of serum samples over time was conducted via robust-trimmed means two-way ANOVA (WRS2::t2way) with mcp2atm used for multiple comparisons, or two-way ANOVA with Tukey’s post-hoc test. Liver weights were analyzed via one-way ANOVA (24 hours, seven days 0.5mg/kg; seven days, 14 days hTYMP dose response) with Tukey’s post-hoc test or unpaired t-tests (14, 21, 28 days 0.5mg/kg). Alpha level for significant differences for all tests was < 0.05.

## Acknowledgements

We are grateful to The Lily Foundation for making this work possible. We thank all lab members for helpful discussions, Z. Golder for technical support and advice, and R. Horvath, P.F. Chinnery, K. Dresser and H. Biggs for continuous support and interest. The authors acknowledge the facilities and the scientific and technical assistance of the Queensland Node of Metabolomics and Proteomics Australia (Q-MAP), BASE and National Biologics Facility (NBF) at The University of Queensland and the Anne McLaren animal facility at the University of Cambridge. Q-MAP is supported by Bioplatforms Australia. BASE and National Biologics Facility (www.nationalbiologicsfacility.com) are supported by Therapeutic Innovation Australia (TIA). Both Bioplatforms Australia and TIA are supported by the Australian Government through the National Collaborative Research Infrastructure Strategy (NCRIS) program. For the purpose of open access, the authors have applied a Creative Commons Attribution (CC BY) license to any Author Accepted Manuscript version arising from this submission.

## Funding

Lily Foundation Research Grant (JvdA, SWC)

MRC Confidence in Concept (JvdA, SWC)

UKRI Medical Research Council (MRC) Mitochondrial Biology Unit MC_UU_00028/8 (JvdA)

Wellcome Clinical Research Career Development Fellowship 219615/Z/19/Z (JvdA)

UKRI BBSRC Responsive Mode Research Grant BB/X00256X/1 (JvdA)

Wellcome Discovery Award 226653/Z/22/Z (JvdA)

UKRI Medical Research Council (MRC) MitoCluster award MC_PC_21046 (JvdA)

Rosetrees Trust PGL23/100048 (JvdA)

LifeArc Centre to Treat Mitochondrial Diseases (LAC-TreatMito) 10748 (JvdA)

Muscular Dystrophy UK (MDUK) (23SI-PRG60-0013) (JvdA)

Australian Research Council FT25010034 (SWC)

Medical Research Future Fund MRFCRI000063 and MRFCRI000089 (SWC)

National Collaborative Research Infrastructure Strategy (NCRIS), Therapeutic Innovation Australia (TIA) (SWC)

Instituto de Salud Carlos III, PI24/00838, funded with ERDF (RM)

Mito Foundation TopUp Scholarship (NLC)

United Mitochondrial Disease Foundation & Mito Foundation Graduate Student Award GS-26-0022 (NLC)

## Author contributions

Conceptualization: JvdA, RM, SWC

Methodology: NLC, JJY, KT, YMK, JR, ZX, AD, TEL

Investigation: NLC, JJY, KT, YMK, SP, JR, ZX, AD, TEL, GN

Funding acquisition: NLC, JvdA, SWC

Project administration: NLC, KT, YMK, JvdA, RM, SWC

Supervision: JvdA, SWC, AQ, RM

Writing – original draft: NLC, JvdA, SWC

Writing – review & editing: all authors

## Data and materials availability

R scripts for statistical analysis are available upon request. Packages used include glmmTMB, lme4, WRS2, ggplot2, emmeans and performance. Raw data for nucleoside measurements, TP activity, Western blots, hematology and serum biochemistry are provided in supplementary materials.

## Supplementary Figures

- Supplementary Figure 1. KO mice treated with mRNA-LNPs show strong liver expression and TP activity.
- Supplementary Figure 2. TP expression and activity are sustained after a single 0.5mg/kg IV dose.
- Supplementary Figure 3. IV administration of *hTYMP*-mRNA-LNPs has no significant effect on blood biochemistry.
- Supplementary Figure 4. IV mRNA-LNPs induce transient peripheral leukopenia.
- Supplementary Figure 5. Acute leukopenia after IV mRNA-LNP delivery is not dose– or genotype-dependent.
- Supplementary Figure 6. SC administration of *hTYMP*-mRNA-LNPs does not induce hepatic expression at low doses.
- Supplementary Figure 7. SC administration of mRNA-LNPs has similar hematological effects to IV administration.
- Supplementary Figure 8. Full Western blot images related to Figure 1.
- Supplementary Figure 9. Full Western blot images related to Figure 2 and Figure 4.
- Supplementary Figure 10. Full Western blot images related to supplementary figures.

## Supplementary Tables

- Supplementary Table 1. Raw data from mass spectrometry of *hTYMP*-mRNA transfected cells. Provided as a separate Excel file.
- Supplementary Table 2. Plasma nucleoside levels. Provided as a separate Excel file.
- Supplementary Table 3. Number of mice per timepoint for monitoring body weight during IV mRNA-LNPs study, related to Figure 3A-D.
- Supplementary Table 4. Serum biochemistry. Provided as a separate Excel file.
- Supplementary Table 5. Whole blood hematology cell counts. Provided as a separate Excel file.
- Supplementary Table 6. Primary and secondary antibodies.

