## Supplementary Figures and Tables S3 and S6 for "mRNA-LNP therapy restores systemic nucleoside imbalance in a mitochondrial DNA depletion syndrome"

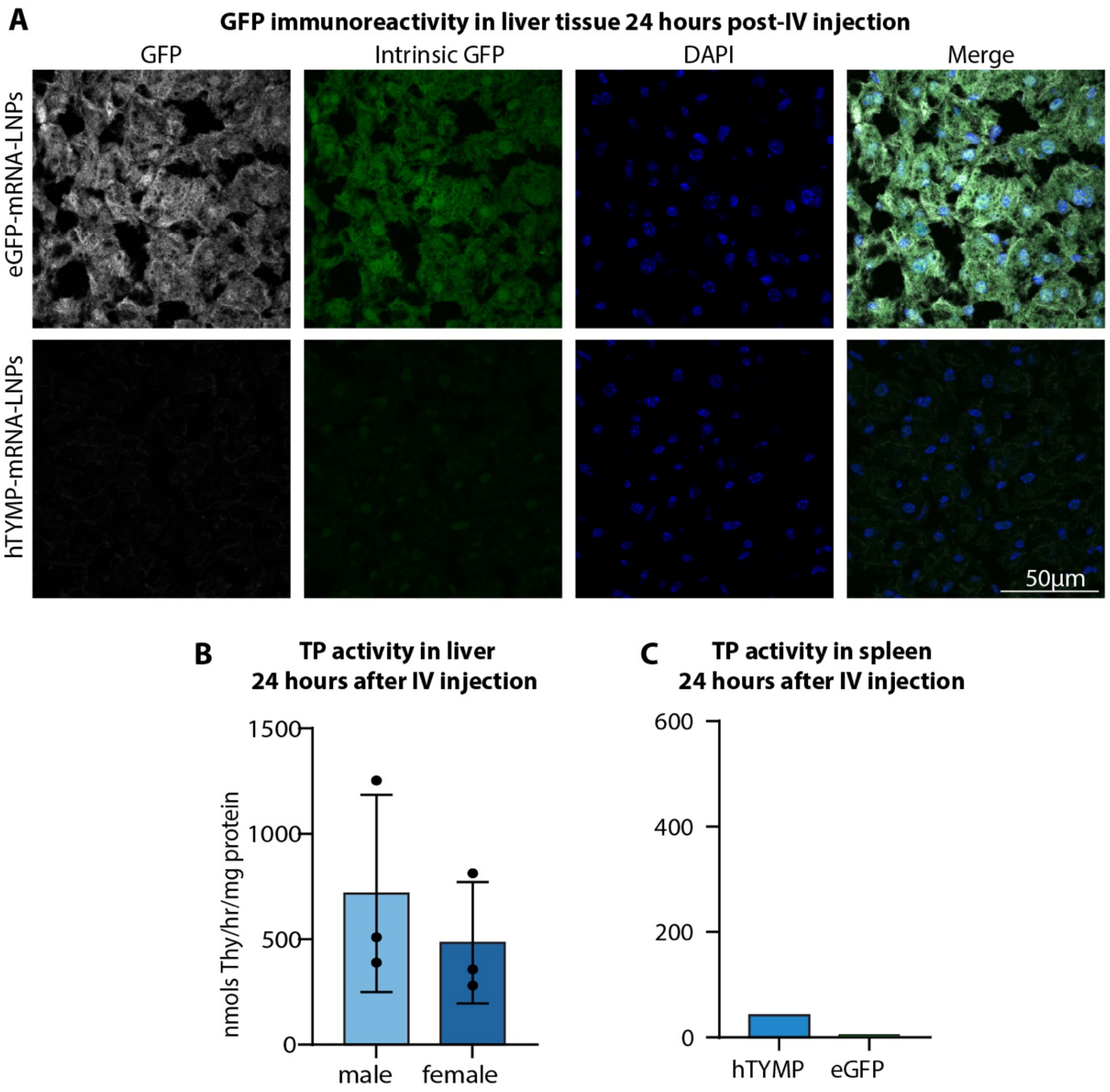


**Supplementary Figure 1. KO mice treated with mRNA-LNPs show strong liver expression and TP activity. (A)** GFP immunostaining (grey), fluorescence (green) and DAPI nuclear stain (blue) in *eGFP*- (top) or *hTYMP*- (bottom) mRNA-LNP treated MNGIE mice 24 hours after IV injection. **(B)** TP enzyme activity in liver of male and female *hTYMP*-mRNA-LNP treated MNGIE mice. Each dot represents one mouse (n = 3). **(C)** TP activity in spleen 24 hours after *eGFP*- or *hTYMP*-mRNA-LNP IV injection in MNGIE mice (n = 1).

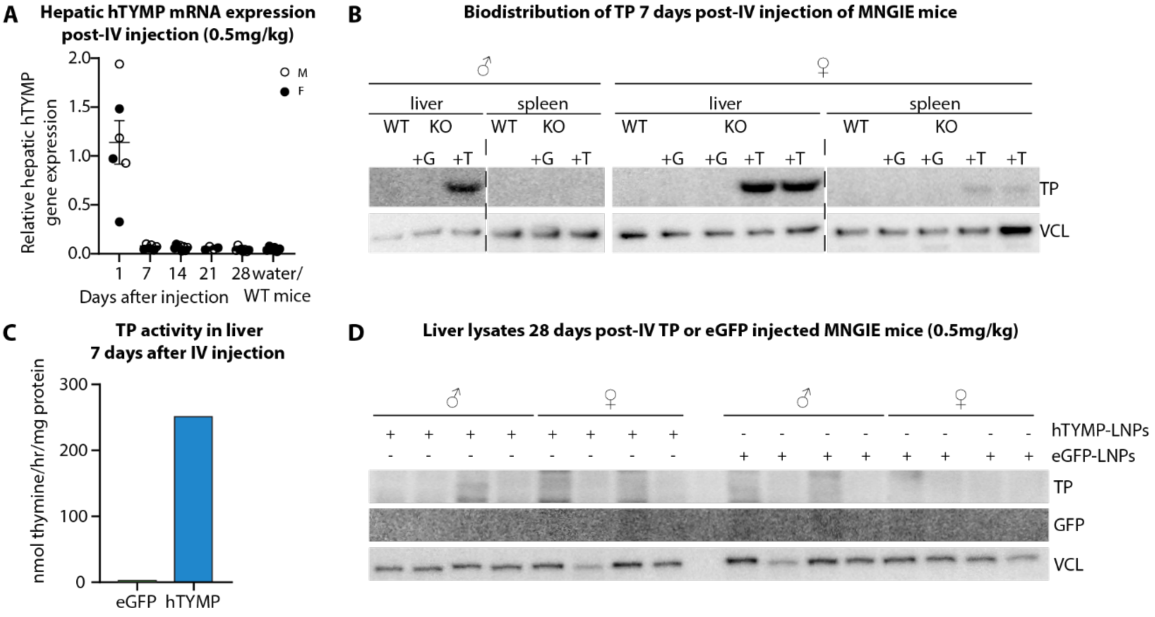


**Supplementary Figure 2. TP expression and activity are sustained after a single 0.5mg/kg IV dose. (A)** *hTYMP* mRNA in livers from *hTYMP*-mRNA-LNP injected mice relative to β-actin mRNA. **(B)** TP expression in livers and spleen from male and female WT and MNGIE mice seven days after intravenous injection. +G = *eGFP*-mRNA-LNPs, +T = *hTYMP*-mRNA-LNPs, WT = wild type mice, KO = MNGIE mice. **(C)** TP activity in liver tissue seven days after delivery of *hTYMP*-mRNA-LNPs. **(D)** TP or GFP expression after 28 days in livers of KO mice treated with *hTYMP*- or *eGFP*-mRNA-LNPs.

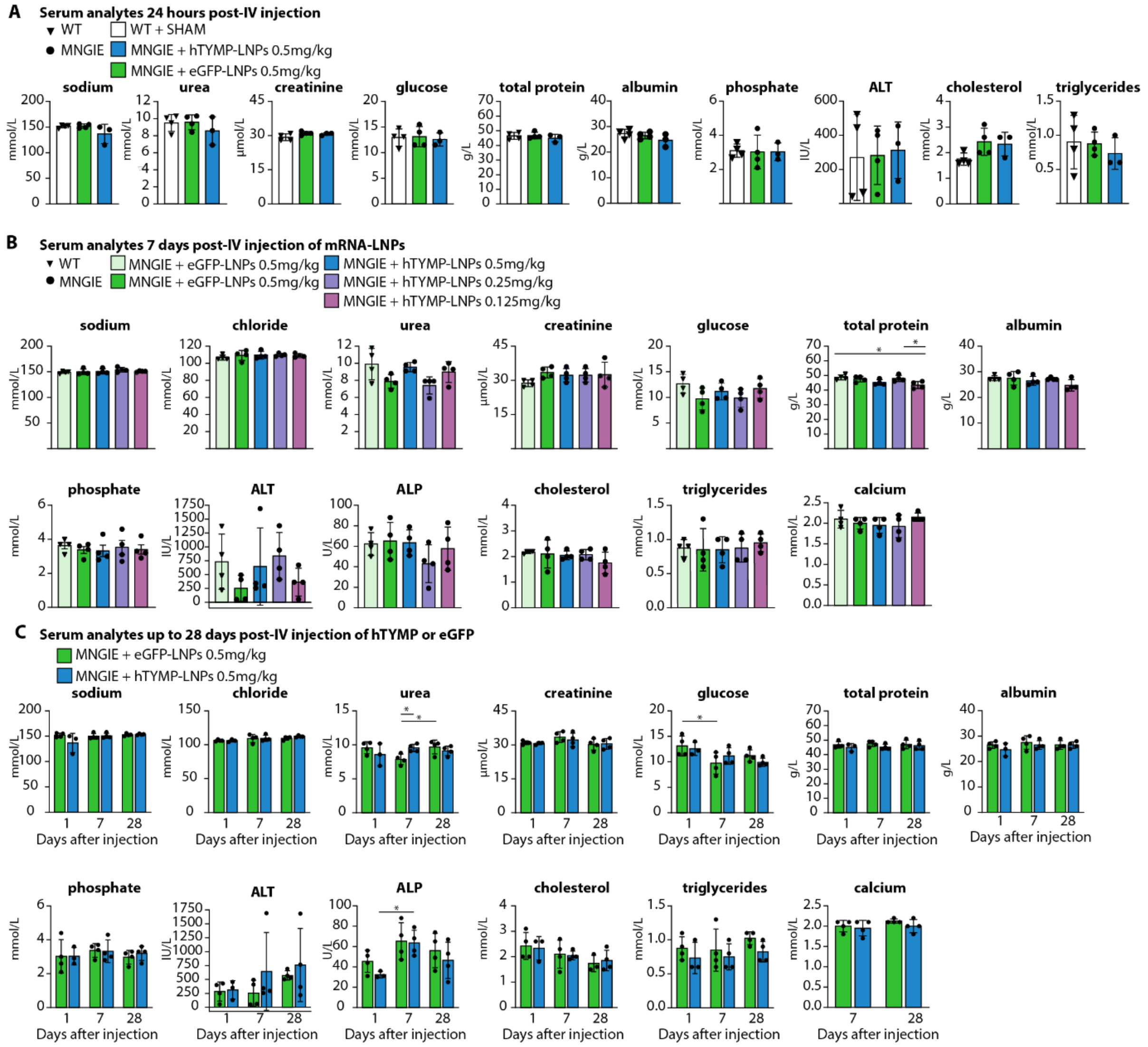


**Supplementary Figure 3. IV administration of *hTYMP*-mRNA-LNPs has no significant effect on blood biochemistry. (A)** The majority of serum analytes show no significant difference when comparing sham-injected wild type mice and mRNA-LNP treated MNGIE mice after 24 hours. One-way ANOVA with Tukey's multiple comparisons or Kruskal-Wallis test. **(B)** Serum biochemistry of wild type (WT) mice and MNGIE mice 7 days after IV administration of mRNA-LNPs. One-way ANOVA with Tukey's multiple comparisons or Kruskal-Wallis test. **(C)** Serum biochemistry values in *eGFP*- or *hTYMP*-mRNA-LNP treated MNGIE mice up to 28 days after IV injection. Two-way ANOVA with Tukey's multiple comparisons or two-way robust trimmed means ANOVA with mcp2way for multiple comparisons. **(A-C)** Column height represents the mean, error bars ± SD. Each dot represents one mouse. * = p ≤ 0.05, ** = p ≤ 0.01, *** = p ≤ 0.001.

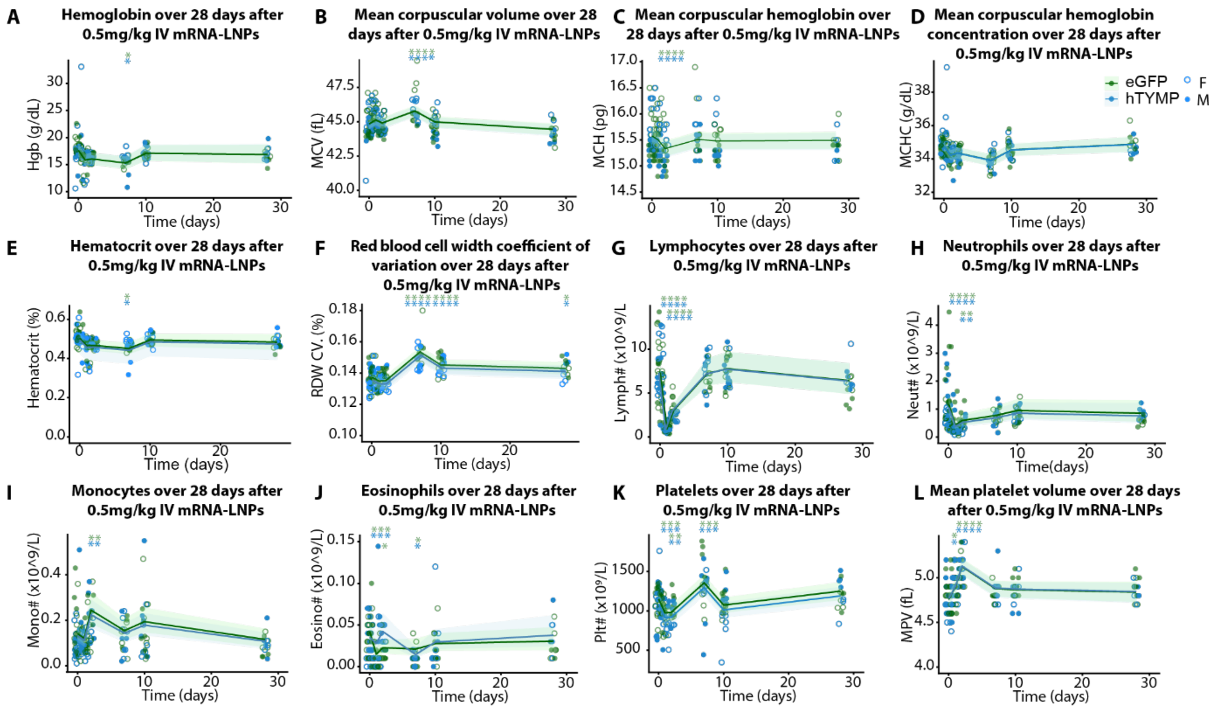


**Supplementary Figure 4. IV mRNA-LNPs induce transient peripheral leukopenia. (A-F)** Red blood cells indices show a significant change from baseline values within the first week aside from mean corpuscular haemoglobin concentration which is unaffected. **(G,H)** Lymphocytes (**G**) and neutrophils **(H)** show a significant decrease from baseline up to 48 hours and recover by 7 days. **(I)** Monocytes show a transient increase 48 hours after mRNA-LNP delivery. (**J**) Eosinophils decrease within the first week and recover by ten days. **(K,L)** Platelets (**K**) decrease and mean platelet volume (**L**) increases significantly up to 48 hours and recover by seven days. Counts were analyzed using generalised linear mixed effects models or linear mixed effects models accounting for mouse and/or sample date as random effects. Lines indicate estimated marginal means and shading shows the 95% confidence intervals. P-values were adjusted for multiple comparisons using the Holm method. Asterisks indicate a significant change compared to the treatment-specific baseline. Each dot represents one mouse. * = p ≤ 0.05, ** = p ≤ 0.01, *** = p ≤ 0.001, **** = p ≤ 0.0001.

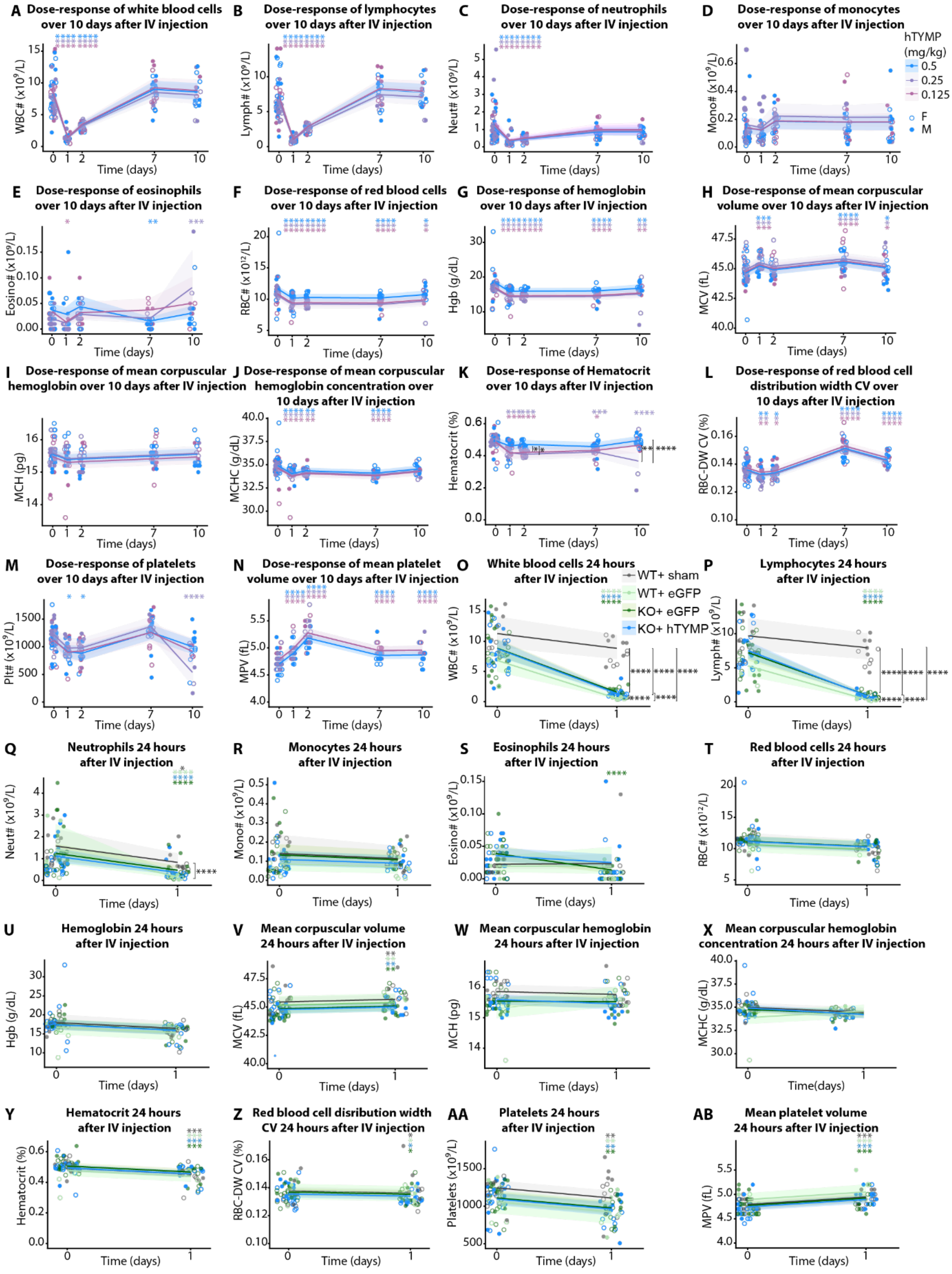


**Supplementary Figure 5. Acute leukopenia after IV mRNA-LNP delivery is not dose- or genotype-dependent. (A-C)** White blood cells, lymphocytes and neutrophils decrease within the first 48 hours regardless of *hTYMP*-mRNA-LNP dose. **(D,E)** Monocytes (**D**) and eosinophils (**E**) remain largely unaffected. **(F-N)** Red blood cell (**F-L**) and platelet (**M,N**) parameters over a ten day period after IV delivery of different doses of *hTYMP*-mRNA-LNPs. **(O-AB)** White blood cell (**O-S**), red blood cell (**T-Z**) and platelet (**AA,AB**) parameters within 24 hours after IV injection of eGFP- or hTYMP-mRNA-LNPs in wild-type (WT) or MNGIE (KO) animals, or in sham injected WT animals. Counts were analyzed using generalised linear mixed effects models or linear mixed effects models accounting for mouse and/or sample date as random effects. Lines indicate estimated marginal means and shading shows the 95% confidence intervals. P-values were adjusted for multiple comparisons using the Holm method. Coloured asterisks indicate a significant change compared to the treatment-specific baseline. Black asterisks show a dose or treatment-specific difference. Each dot represents one mouse. * = $p \leq 0.05$, ** = $p \leq 0.01$, *** = $p \leq 0.001$, **** = $p \leq 0.0001$.

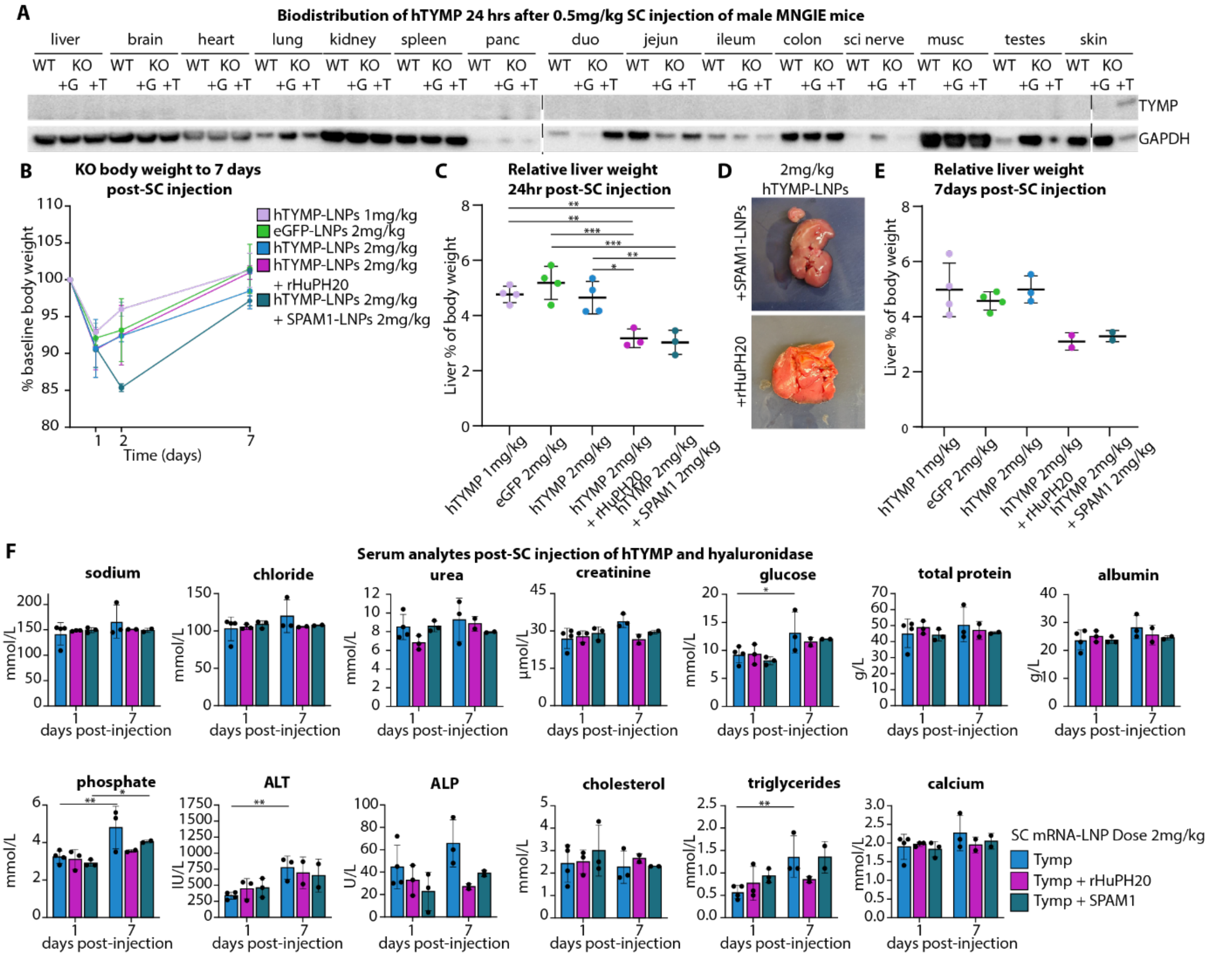


**Supplementary Figure 6. SC administration of *hTYMP*-mRNA-LNPs does not induce hepatic expression at low doses. (A)** TP expression in indicated tissues 24 hours after subcutaneous (SC) injection of 0.5mg/kg *hTYMP*-mRNA-LNPs. KO = MNGIE (*Tymp1*$^{-/-}$ /*Upp1*$^{-/-}$) knockout mice. +G = *eGFP*-mRNA-LNP, +T = *hTYMP*-mRNA-LNP. **(B)** Mean body weight decreases by 10-15% within 48 hours of SC mRNA-LNP delivery but recovers by seven days. **(C-E)** Relative liver weights (**C**,**E**) and appearance (**D**) 24 hours (**C**,**D**) or 7 days (**E**) after administration of mRNA-LNPs with or without hyaluronidase, either as rHuPH20 or *SPAM1*-mRNA-LNPs. **(F)** Serum biochemistry up to seven days in KO mice treated SC with *hTYMP*-mRNA-LNPs with or without hyaluronidase, either as rHuPH20 or *SPAM1*-mRNA-LNPs. Column height represents the mean, error bars show ± SD. One-way ANOVA with Tukey's multiple comparisons **(C)**. Two-way ANOVA with Fisher's least significant difference **(F)**. * = p ≤ 0.05, ** = p ≤ 0.01, *** = p ≤ 0.001. Each dot represents data from one mouse.

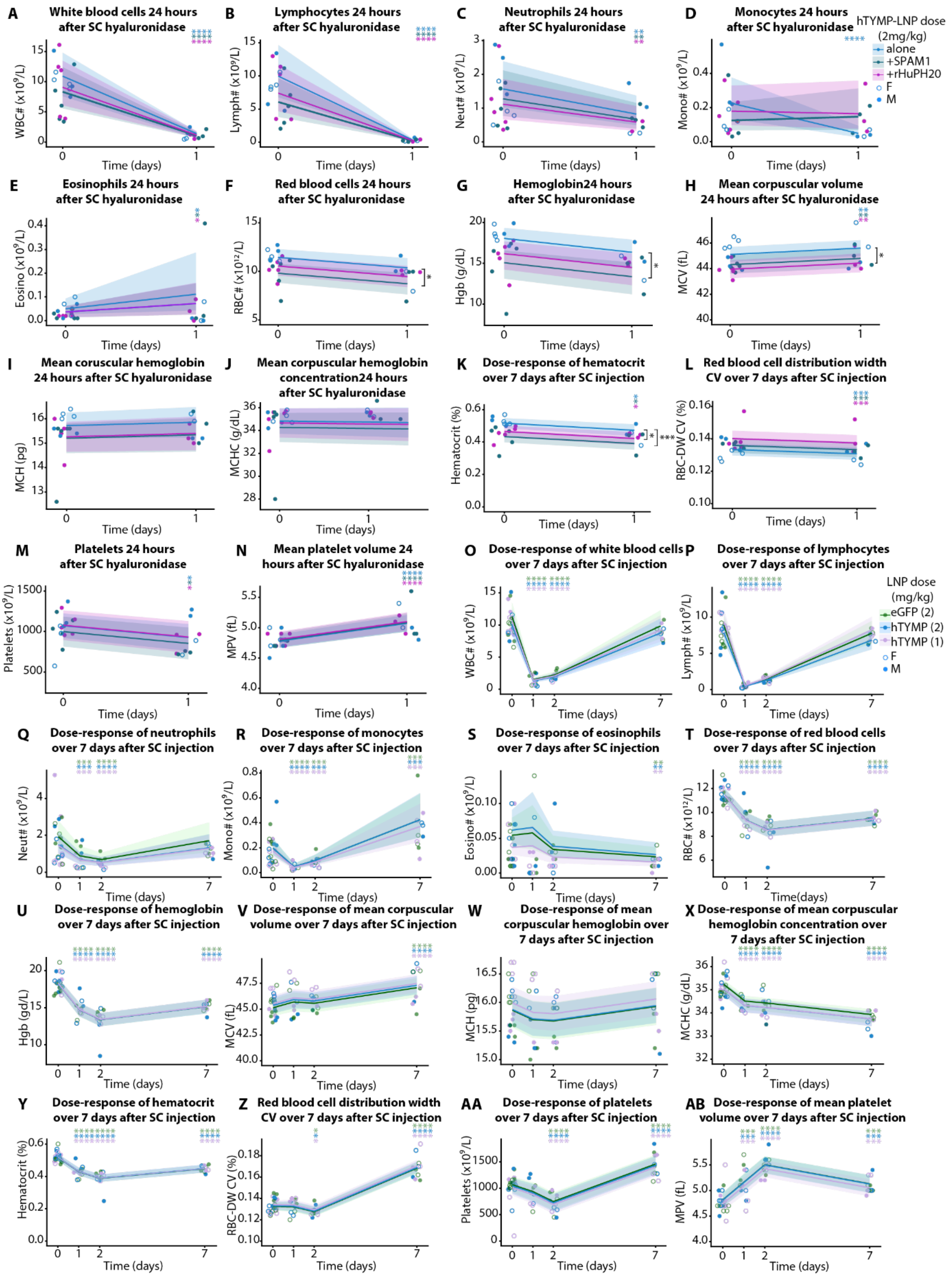


**Supplementary Figure 7. SC administration of mRNA-LNPs has similar hematological effects to IV administration. (A-N)** White blood cell (**A-E**), red blood cell (**F-L**) and platelet

(**M,N**) parameters 24 hours after subcutaneous (SC) mRNA-LNP delivery without (blue) or with hyaluronidase as *SPAM1*-mRNA-LNPs (green) or rHuPH20 (purple).

**(O-AB)** White blood cell (**O-S**), red blood cell (**T-Z**) and platelet (**AA,AB**) parameters within seven days after subcutaneous (SC) mRNA-LNP delivery of 2mg/kg *eGFP*-mRNA-LNPs (green), 2mg/kg *hTYMP*-mRNA-LNPs (blue) or 1mg/kg *hTYMP*-mRNA-LNPs (purple). Counts were analyzed using linear mixed effects or generalised linear mixed effects models with multiple comparisons adjusted via Tukey's or Holm methods. Coloured asterisks mark a significant change compared to the treatment-specific baseline. Black asterisks show a treatment-specific difference. Each dot represents one mouse. * = $p \leq 0.05$, ** = $p \leq 0.01$, *** = $p \leq 0.001$, **** = $p \leq 0.0001$.

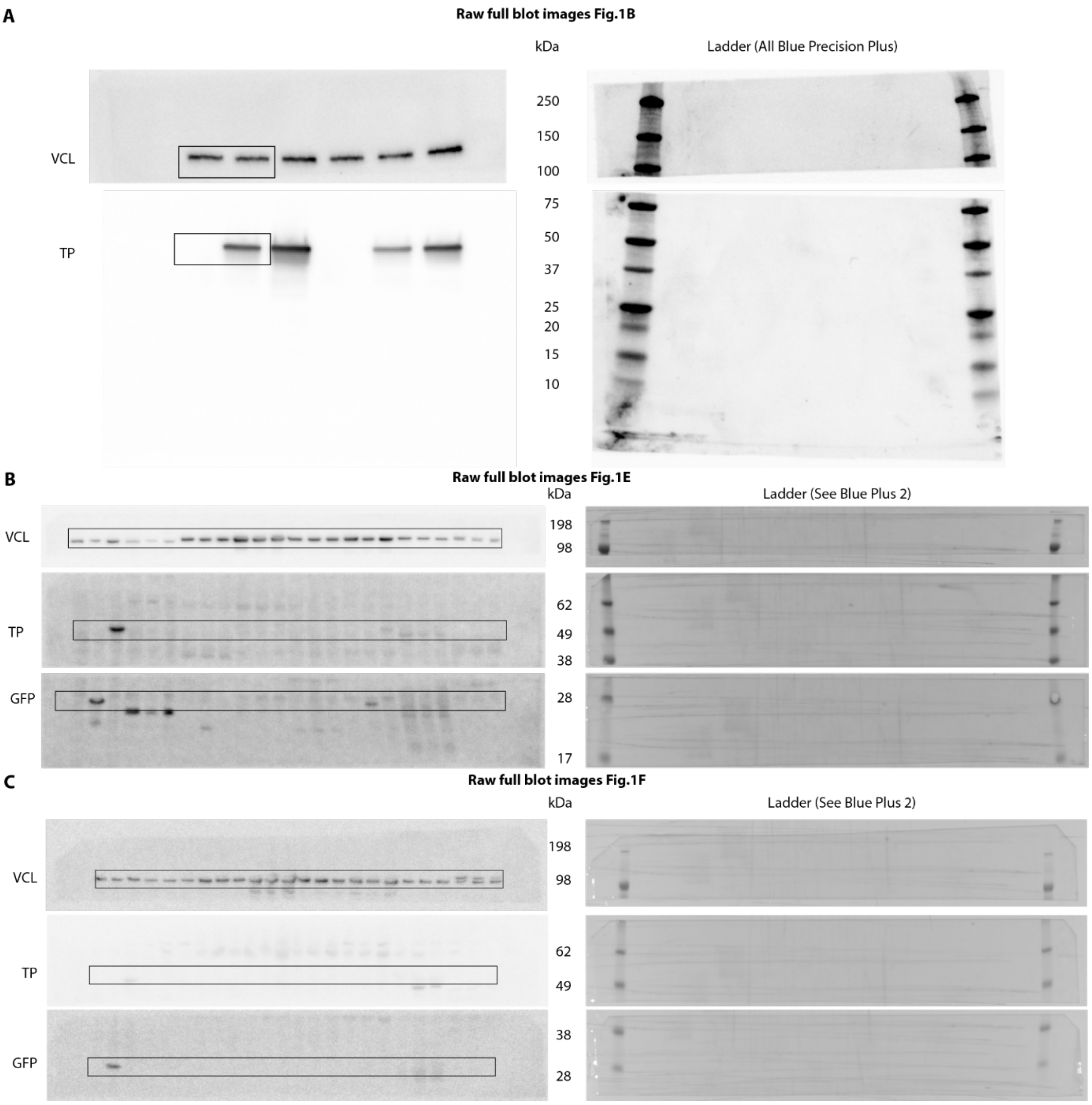


**Supplementary Figure 8.** Full Western blot images related to Figure 1.

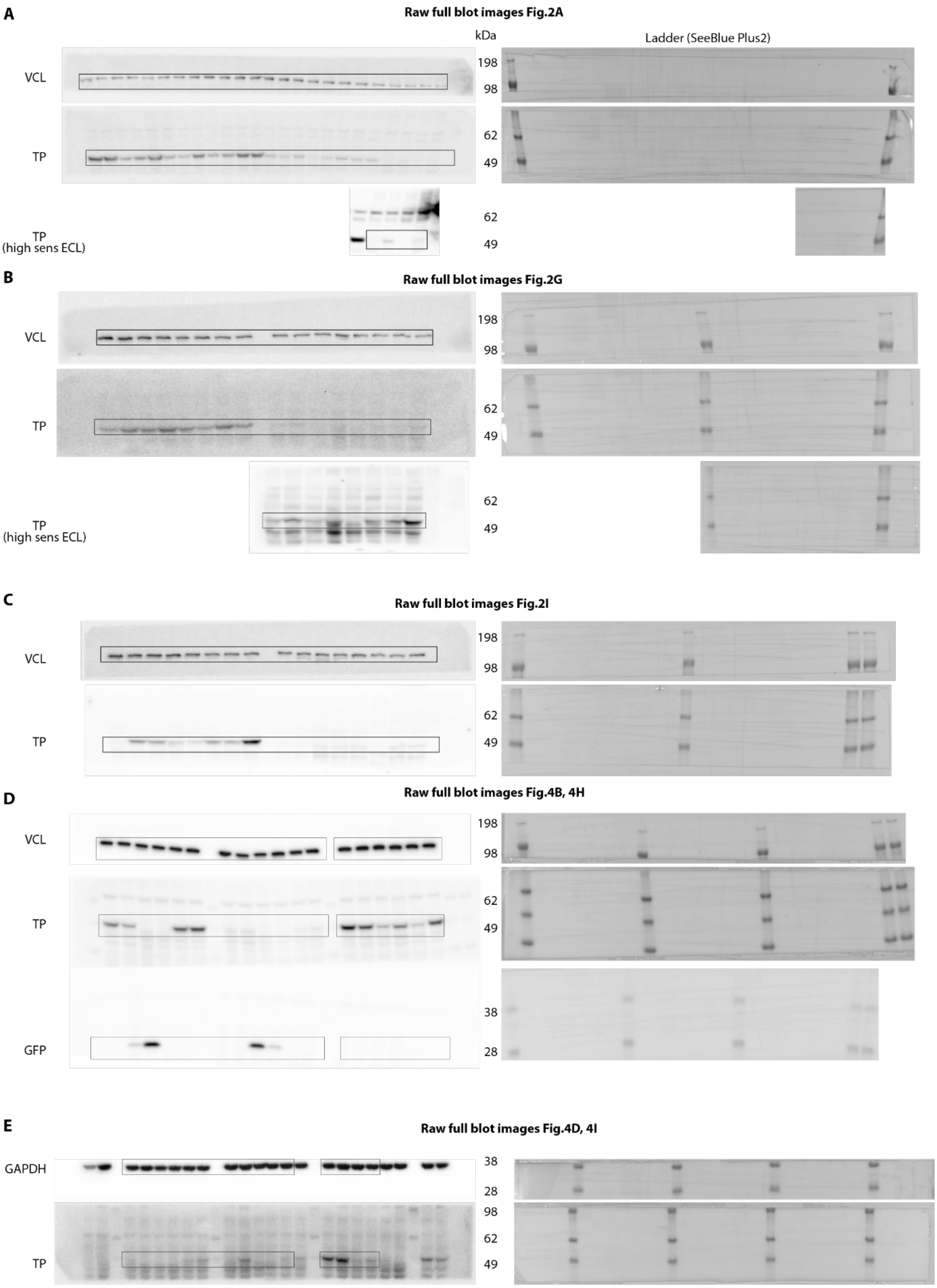


**Supplementary Figure 9.** Full Western blot images related to Figure 2 and Figure 4.

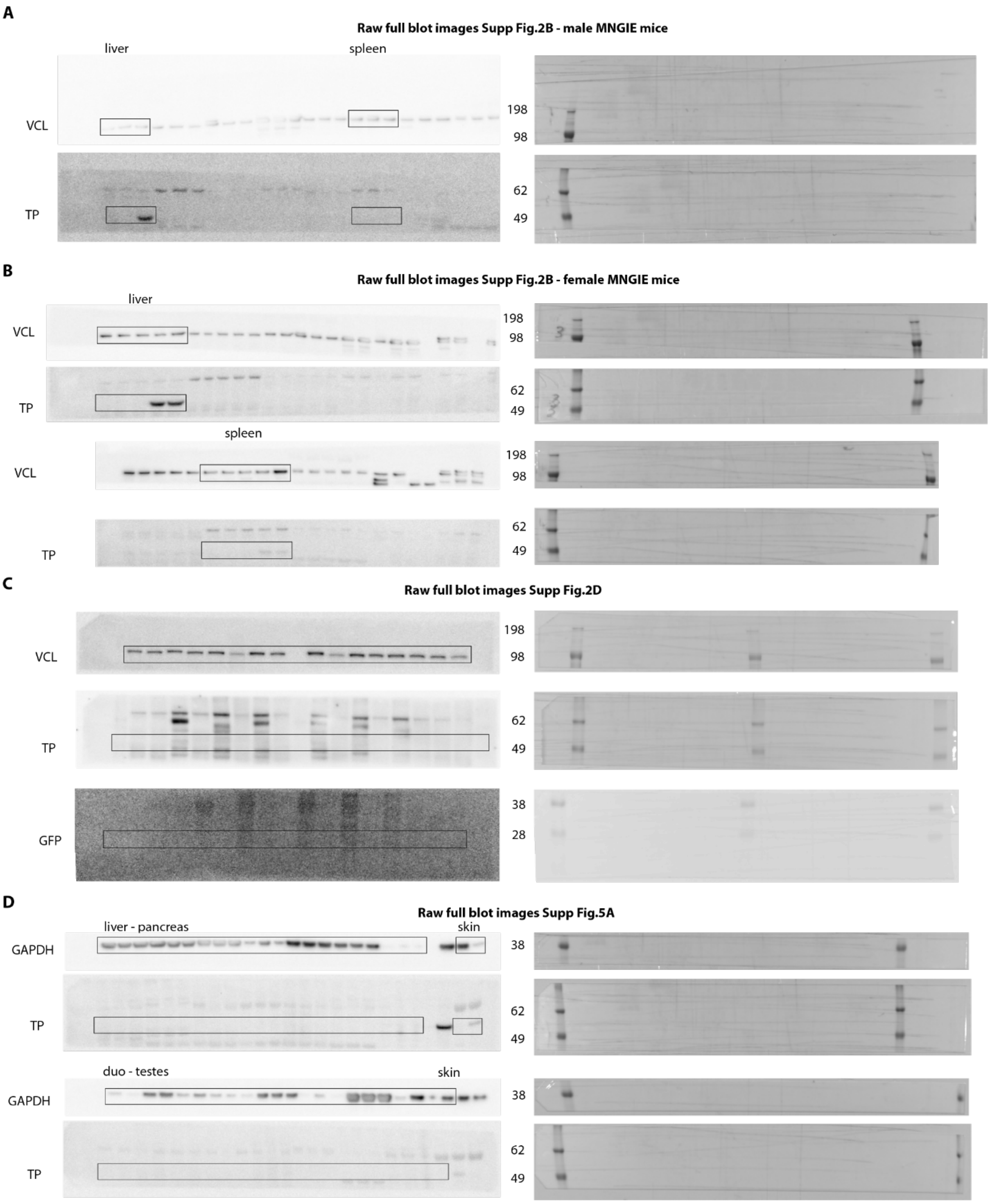


**Supplementary Figure 10.** Full Western blot images related to supplementary figures.

**Supplementary Tables**

**Supplementary Table 1.** Raw data from mass spectrometry of *hTYMP*-mRNA transfected cells. Provided as a separate Excel file.

**Supplementary Table 2.** Plasma nucleoside levels. Provided as a separate Excel file.

**Supplementary Table 3.** Number of mice per timepoint for monitoring body weight during IV mRNA-LNPs study, related to Figure 3A-D.

| | WT mice | | MNGIE (KO) mice | | | |
|---|---|---|---|---|---|---|
| Treatment | Sham | eGFP-mRNA-LNPs | eGFP-mRNA-LNPs | hTYMP-mRNA-LNPs | | |
| Timepoint (days) | | 0.5mg/kg | 0.5mg/kg | 0.5mg/kg | 0.25mg/kg | 0.125mg/kg |
| 1 | 12 | 6 | 20 | 20 | 8 | 8 |
| 2 | - | 3 | 16 | 16 | 8 | 10 |
| 7 | - | 6 | 14 | 14 | 8 | 8 |
| 10 | - | - | 12 | 12 | 4 | 4 |
| 14 | - | - | 8 | 8 | 8 | 8 |
| 21 | - | - | 4 | 4 | - | - |
| 28 | - | - | 8 | 8 | - | - |

**Supplementary Table 4.** Serum biochemistry. Provided as a separate Excel file.

**Supplementary Table 5.** Hematological cell counts. Provided as a separate Excel file.

**Supplementary Table 6.** Primary and secondary antibodies

| **Antibody** | **Catalogue #** | **Supplier** | **Application** | **Working concentration** |
|---|---|---|---|---|
| Mouse anti-Vinculin | V4505 | Merck | Western blot | 15-55µg/mL |
| Rabbit anti-TYMP | PA5-81917 | Invitrogen | Western blot | 0.2µg/mL |
| Rabbit anti-GAPDH | 5174 | Cell Signalling Technologies | Western blot | 34ng/mL |
| Rabbit anti-GFP | 6556 | Abcam | Western blot | 0.5µg/mL |
| Swine anti-rabbit-HRP | P0217 | Dako | Western blot | 1.3µg/mL |
| Rabbit anti-mouse-HRP | P0260 | Dako | Western blot | 1.3µg/mL |
| Rabbit anti-TYMP N terminal | 180783 | Abcam | Immunofluorescence | 7.5µg/mL |
| Chicken anti-GFP | 13970 | Abcam | Immunofluorescence | 20µg/mL |
| Alexa Fluor 546 anti-chicken | A11040 | Thermo Fisher | Immunofluorescence | 4µg/mL |
| Alexa Fluor 555 anti-rabbit | A32794 | Thermo Fisher | Immunofluorescence | 4µg/mL |
